# Migratory dermal γδ T cells determine the balance between lung immunity and tissue damage during *Nippostrongylus brasiliensis* infection

**DOI:** 10.1101/2025.01.07.631663

**Authors:** Pedro H. Papotto, James E. Parkinson, Brian H.K. Chan, I-Hsuan Lin, Georgia E. Baldwin, Faith Denny, Lili Zhang, Rebecca J. Dodd, Stella T. Pearson, Natalie Fischhaber, Joanne E. Konkel, David R. Withers, Matthew R. Hepworth, Judith E. Allen

## Abstract

Many helminth parasites migrate through multiple host organs during infection but how immune responses are relayed across these tissues is still poorly understood. To investigate the cellular and molecular aspects of inter-tissue communication during infection we established a percutaneous infection model with the tissue-migrating nematode *Nippostrongylus brasiliensis*. High-dimension profiling of the initial cutaneous immune response revealed that dermal γδ T cells become activated via an IL-1R-dependent mechanism, engage cell motility-associated transcriptional pathways and leave the skin after parasite invasion. Chemical and genetic inhibition of leukocyte migration prevents the accumulation of IL-17-producing γδ T cells in the lungs. Notably, bypassing the skin phase of infection, and therefore preventing dermal γδ T cell migration, dampens the increase in early IL-17 production in the lungs, and, instead, leads to enhanced IFN-γ responses together with increased lung damage. Collectively, our data highlights a critical skin–lung axis regulating host-parasite interactions and safeguarding lung health.

## Introduction

The mammalian immune system evolved to ensure that upon pathogen invasion organ-specific responses are employed appropriately to promote host survival(1, 2). Achieving a balance that leads to pathogen elimination, whilst maintaining tissue integrity often requires the co-ordination of conflicting immune responses within particular tissues(3). Reaching this equilibrium is even more complex when we consider pathogens that migrate across different organs. As a result, our understanding of how immunity is orchestrated at the whole-organism level is still lacking.

Large multicellular parasites such as helminths are usually associated with type-2 immunity(4). Given the size of these pathogens and their (usually long) life cycles within the host, it is believed that the ultimate goal of type-2 immune responses is to prevent parasite migration through host tissues and to drive worm expulsion whilst keeping collateral damage to a minimum(5). Hence, type-2 immunity promotes different degrees of tissue remodelling(6). The rodent parasite *Nippostrongylus brasiliensis* has been widely used to study the mechanisms regulating innate and adaptive type-2 immunity(7). The *N. brasiliensis* experimental model mimics key aspects of human hookworm infection, with larvae entering the host through the skin, and reaching the lung 18 to 48 hours post-infection (p.i.), where they moult before migrating to the small intestine(7). There, they become sexually mature and start releasing eggs before being expelled from the gut by 6 to 8 days p.i.(7). The importance of type-2 immunity in controlling *N. brasiliensis* infection is well-documented(4, 8). Surprisingly, work from our group has shown that in the lungs *N. brasiliensis* initially induces IL-17A production by γδ T cells, which in turn is critical for the generation of subsequent type-2 immunity(9, 10). At the same time, IL-17A promotes neutrophil recruitment, contributing to increased tissue damage(11, 12), while also enhancing IL-33 release and activation of ILC2s(11). In addition, IL-17A further enables a type-2-promoting environment by suppressing Interferon-γ (IFN-γ) production by γδ T cells and other innate-like lymphocytes(9), through mechanisms still not fully understood.

Despite early evidence of γδ T cell responses to helminth infections(13), the mechanisms behind their activation by these parasites are poorly defined. In contrast to conventional αβ CD4^+^ and CD8^+^ T cells, most murine γδ T cells acquire their effector functions during thymic development, reaching peripheral tissues readily capable of producing IL-17A or IFN-γ(14, 15). This functional dichotomy in murine γδ T cells is tightly linked to the variable γ (Vγ) chain of their T cell receptor (TCR)(15). Notably, γδ T cells in the dermal layer of the skin are exclusively IL-17-producing γδ (γδ17) T cells, bearing Vγ4 and Vγ6 TCRs(15, 16). Dermal γδ T cells are a motile population that exhibits a slow but steady migration pattern to the skin-draining lymph nodes(16, 17). However, whether these cells have the ability to migrate to other non-lymphoid organs during inflammation – akin to intestinal γδ17 T cells(18, 19) – is yet to be determined. Thus, we hypothesised that activation of skin γδ T cells by *N. brasiliensis* has consequences beyond their tissue of residency, directly regulating the lung immune response to the parasite.

Here we optimised a percutaneous (p.c.) *N. brasiliensis* infection model mimicking the natural steps of parasite invasion and migration through the skin. This allowed us to generate a comprehensive analysis of the early immune events taking place in the skin from 6 to 24 hours after larval invasion. *N. brasiliensis* larvae induced the activation of dermal Vγ6 γδ T cells via an IL-1R-dependent mechanism, which, in turn, upregulated pathways related to cell migration and relocated to the lungs, where they were responsible for the increase in γδ17 T cells within the tissue. Dermal γδ T cell migration to the lungs was dependent on CCR2 and S1PR; suppression of these pathways affected both the lung immune response to the parasite and tissue damage control. Finally, bypassing the skin phase of the infection through intravenous (i.v.) injection of *N. brasiliensis* larvae prevented dermal γδ17 T cell migration, whilst inducing the expansion of lung IFN-γ producing γδ (γδ1) T cells and increased tissue damage.

## Results

### *Percutaneous* N. brasiliensis *infection recapitulates the major hallmarks of the subcutaneous injection model*

Standard laboratory practice of inducing *N. brasiliensis* infection involves subcutaneous (s.c.) injection of larvae. While practical, it fails to recapitulate natural events in the skin that may affect subsequent immune events in distal organs. We thus needed a model that elicits immune responses along all the strata of the skin and avoids needle-induced effects but also generates a consistent and reproducible infection in the lung and gut. To achieve this, we adapted a protocol from the 1970’s(20) and placed a moist filter paper containing 500 N. brasiliensis L3s on the ear of an anaesthetised mouse for 30 minutes **(Fig. 1a)**. This protocol ensures larvae penetrate through the skin, and we confirmed that the parasites could be found near the base of the ear, up until 12h p.i. **(Fig. 1b)**; at 24h p.i. tissue damage was evident, but larvae were no longer visible in the skin **(Fig. 1b)**. NanoString analysis of whole-skin tissue indicated that transcriptional changes were evident by 6h p.i., with the biggest differences observed at 24h p.i. **(Extended Data Fig. 1a and b)**. Pathway analysis showed that differentially upregulated genes at 24h p.i. related to cytokine signalling, response to LPS and type-I IFN production, supporting published findings using non-viable parasites injected intradermally(21, 22) **(Extended Data Fig. 1c)**. Differentially downregulated genes, on the other hand, were mostly associated with pathways regulating cell-cell adhesion **(Extended Data Fig. 1d)**. An increase in skin-draining lymph node cellularity **(Extended Data Fig. 1e)** provided further evidence of immune activation in the skin. Importantly, the p.c. route led to infection kinetics akin to the s.c. model **(Extended Data Fig. 1f)** with the parasite being found in the lungs between days 1 and 2 p.i., then in the intestines between days 2 and 5 p.i., being mostly cleared after day 6 p.i. **(Fig. 1c)**. Type-2 immunity also followed the kinetics of s.c. infections(23), with p.c.-infected mice showing an increase in IL-5^+^ ILC2s as early as day 1 p.i., and IL-13^+^ ILC2s from days 5 to 12 **(Extended Data Fig. 1g)**. We found that IL-5 **(Extended Data Fig. 1h)** and IL-13 **(Extended Data Fig. 1i)** production by CD4^+^ T cells peaked at day 12 p.i., with eosinophil recruitment following the same pattern **(Extended Data Fig. 1j)**.

**Fig. 1.**
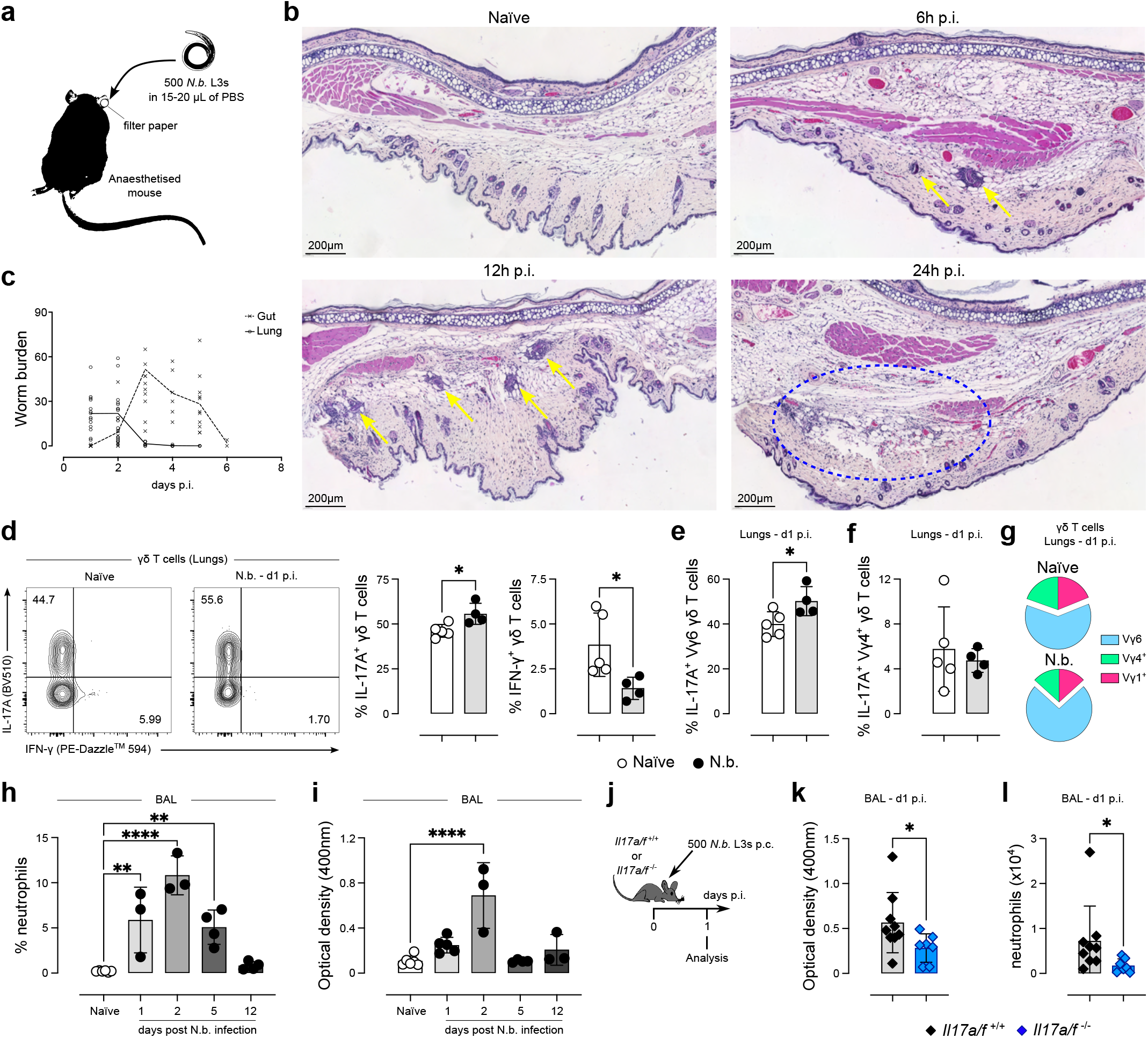
*N. brasiliensis* p.c. infection recapitulates the hallmarks of the s.c. route of infection. **a**, Experimental procedure for *N. brasiliensis* p.c. infection. 500 L3s were added to a piece of filter paper and placed on the ear of an anaesthetised mouse for 30 minutes. **b**, Representative H&E staining micrography of ear sections from naïve (top left) and *N. brasiliensis*-infected C57BL/6J mice at 6h (top right), 12h (bottom left) and 24h p.i. (bottom right); yellow arrows indicate *N. brasiliensis* larvae, and the blue circle shows an inflammatory focus. **c**, worm burden measured in the lungs (full line) and small intestines (dashed line) throughout the course of *N. brasiliensis* p.c. infection (*n* = 3-16 mice per organ and time point; with the exception of days 4 and 6 p.i., which were done once, every timepoint has been repeated 3 times). **d**, Flow cytometry analysis of intracellular IL-17A and IFN-γ in γδ T cells isolated from the lungs of naïve and *N. brasiliensis*-infected C57BL/6J mice and stimulated ex vivo for 4h. Summary graph showing the percentages of IL-17A^+^ and IFN-γ^+^ cells within the γδ T cell population (*n* = 4-5 mice per group; representative of at least 3 independent experiments). **e, f**, Frequencies of IL-17A^+^ cells within the Vγ6 γδ T cell population (defined as Vγ1^-^Vγ4^-^) **(e)** and the Vγ4^+^ γδ T cell population **(f)** in the lungs of naïve and *N. brasiliensis*-infected C57BL/6J mice (*n* = 4-5 mice per group; representative of at least 3 independent experiments). **g**, Pie charts depicting the Vγ chain usage among lung γδ T cell in naïve and *N. brasiliensis*-infected C57BL/6J mice (*n* = 4-5 mice per group; representative of at least 3 independent experiments). **h**, Frequencies of neutrophils within total leukocytes in the BAL throughout the course of *N. brasiliensis* p.c. infection (*n* = 3-6 mice per group; every time point has been repeated individually at least 3 times). **i**, Optical density of the BAL of naïve and *N. brasiliensis*-infected C57BL/6J mice measured at 400nm to quantify lung bleeding (*n* = 3-6 mice per group; every time point has been repeated individually at least 3 times). **j**, *Il17a/f* ^-/-^ and their WT littermate controls were infected as depicted in **(a)** and analysed at da 1 p.i. **k**, Optical density of the BAL of *N. brasiliensis*-infected *Il17a/f* ^-/-^ and *Il17a/f* ^+/+^ mice measured at 400nm to quantify lung bleeding (*n* = 7-9 mice per group). **l**, neutrophil numbers in the BAL of N. brasiliensis-infected *Il17a/f* ^-/-^ and *Il17a/f* ^+/+^ mice (*n* = 7-9 mice per group). **c-k**, Error bars represent mean±SD. Normality of the samples was assessed with Shapiro-Wilk normality test; statistical analysis was then performed using Student’s t test **(d, e)**, Mann-Whitney test **(k**,**l)**, or one-way ANOVA test (e, f). * *p* < 0.05; ** *p* < 0.01; **** *p* < 0.0001

Critical for this study, *N. brasiliensis* p.c. infection induced an expansion in the frequency of lung γδ17 T cells at day 1 p.i. (with cell numbers going up by day 2 p.i.; **Extended Data Fig. 1k**) and a concurrent decrease in IFN-γ-producing γδ (γδ1) T cells **(Fig. 1d)**, corroborating published data using s.c. infection(9, 10). Of note, the increase in IL-17A production was derived from Vγ6, but not Vγ4^+^, γδ T cells **(Fig. 1e, f)**, which were also enriched in the lungs at the population level upon *N. brasiliensis* infection **(Fig. 1g)**. Accordingly, we found neutrophil levels to be increased in the bronchoalveolar lavage (BAL) as early as day 1 p.i., remaining elevated until day 5 p.i., and returning to naïve levels at day 12 p.i. **(Fig. 1h)**. Lung damage was characterised by evidence of haemorrhage in the BAL, which followed the pattern of neutrophil kinetics **(Fig. 1i)**, mirroring the findings from the s.c. model(24) **(Extended Data Fig. 1l, m)**. Importantly, using our *N. brasiliensis* p.c. infection model in *Il17a/f* ^+/+^ and *Il17a/f* ^-/-^ mice **(Fig. 1j)** we confirmed published data by us and others(9, 10, 12) indicating that IL-17 deficiency results in decreased BAL haemorrhage and neutrophil recruitment **(Fig. 1k, l)**, together with the inability to regulate IFN-γ-related transcriptional pathways **(Extended Data Fig. 1n)**. Overall, our p.c. *N. brasiliensis* infection model recapitulates the hallmarks of the pulmonary response against migrating larvae, making it a useful tool in the study of skin – lung communication.

### Bypassing the skin phase of infection results in increased IFN-γ production and lung haemorrhage

To disentangle the contribution of the skin phase of infection from subsequent lung immunity we compared p.c.-infected mice with animals that were infected intravenously (i.v.) with *N. brasiliensis* larvae **(Fig. 2a)**. On a tissue-wide level, NanoString analysis indicated that p.c. and i.v. infections induce distinct transcriptional changes in the lungs **(Fig. 2b)**, with p.c.-infected mice showing increased expression of genes related to neutrophil recruitment (such as *Sele* and *Cxcl2*) and their i.v. counterparts exhibiting an increase in the expression type-1 effector genes, such as *Ccl5* and *Klrk1* **(Fig. 2c)**. Of note, these discrete transcriptional changes were not due to differences in infection efficacy, as the two routes of infection led to a comparable worm burden in the lungs at days 1 and 2 p.i. **(Extended Data Fig. 2a)**. As expected from the previous flow cytometry data **(Fig. 1d)**, *N. brasiliensis* p.c.-infected mice showed an enrichment in pathways related to IL-17 signalling **(Fig. 2d)**, which was something not observed in i.v.-infected animals that in turn displayed an enrichment in pathways related to wound healing **(Fig. 2e)**. Accordingly, *N. brasiliensis* i.v. infections did not induce an increase in lung γδ17 T cells, in contrast to what we observed in p.c. infections **(Fig. 2f)**. Moreover, i.v.-infected mice also did not display an increase in lung Vγ6 γδ T cells, in a clear contrast to p.c.-infected mice **(Fig. 2g and Extended Data Fig. 2b)**. Importantly, intravenous CD45 labelling(25) indicated that *N. brasiliensis* p.c. (but not i.v.) infection induces the increase of total γδ T cells and Vγ6 γδ T cells within the pulmonary tissue and not within the circulating population **(Extended Data Fig. 2c, d)**. Instead, i.v.-infected mice showed an increase in the frequency of IFN-γ-producing γδ (γδ1) T cells in the lungs when compared to naïve and p.c.-infected animals **(Fig. 2f)**. In addition, only *N. brasiliensis* i.v. infection led to enhanced type-2 cytokine production, with increased levels of IL-13, IL-9 and IL-5 **(Extended Data Fig. 2e)**. As expected from their failure to induce γδ17 T cell responses, i.v.-infected mice also failed to recruit neutrophils to the lungs **(Fig. 2h)**. To our surprise, however, despite not inducing a neutrophilic response in the lungs, bypass of the skin resulted in increased lung haemorrhage **(Fig. 2i and Extended Data Fig. 2f)**. An increase of neutrophils, eosinophils, T cells and B cells was observed in the BAL of i.v.-infected mice but not in p.c.-infected mice **(Extended Data Fig. 2g)**, possibly reflecting the differential haemorrhagic responses in these mice **(Fig 2h)**, and supported by increased staining with intravenous anti-CD45 **(Extended Data Fig. 2h)**. Because IFN-γ has been shown to promote lung bleeding during viral infection(26), we infected *Ifngr*^-/-^ mice and their *Ifngr*^+/+^ littermates with *N. brasiliensis* i.v. **(Fig. 2j)**. *Ifngr*^-/-^ mice exhibited a decrease in lung haemorrhage, when compared to wild-type (WT) controls **(Fig. 2k and Extended Fig. 2i)**. Thus, our results indicate that bypassing the skin phase of *N. brasiliensis* infection fails to induce an increase in γδ17 T cells or associated neutrophil recruitment, and instead induces IFN-γ production that drives increased lung bleeding.

**Fig. 2.**
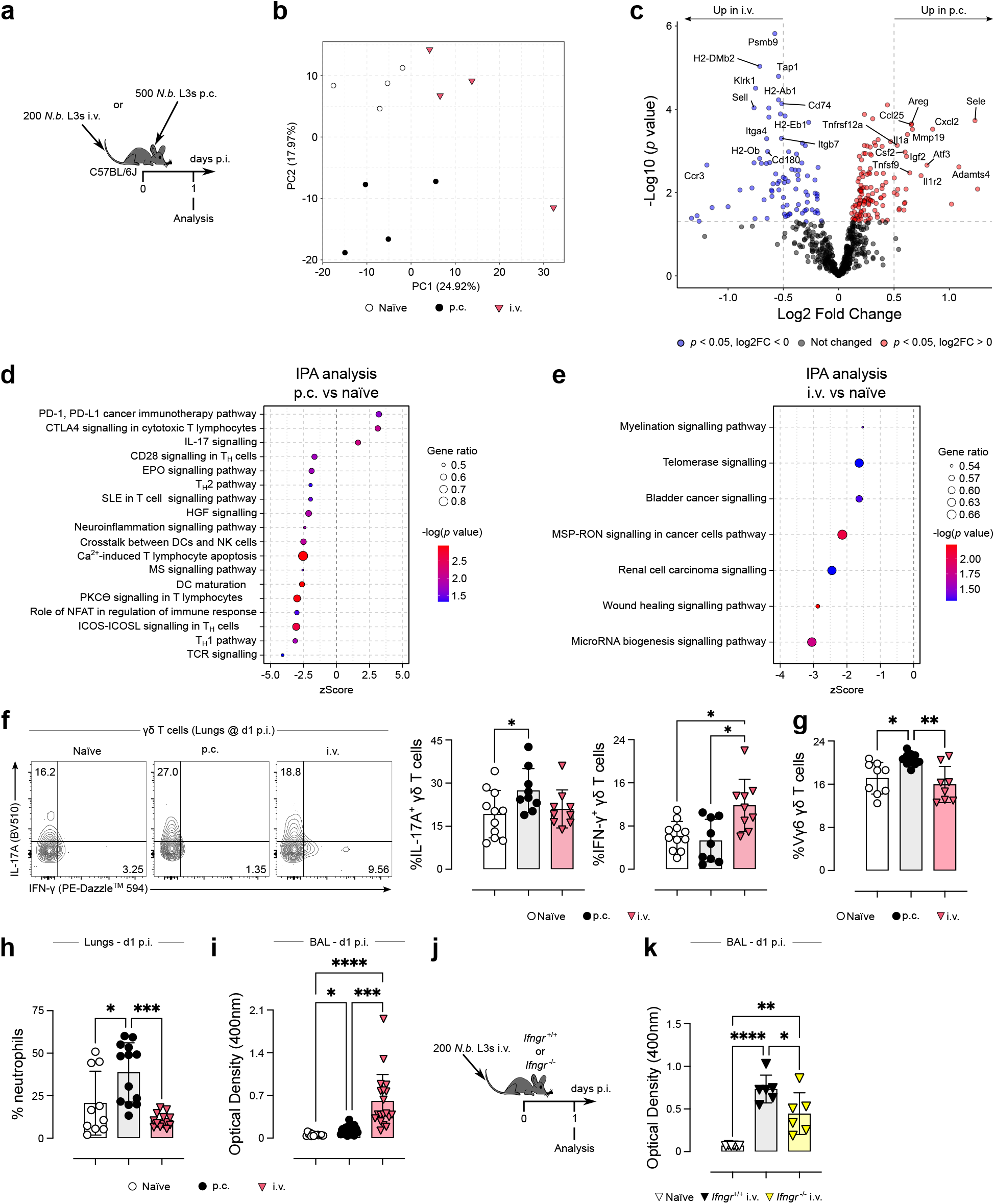
Bypassing the skin phase of infection precludes the induction of lung type-17 responses and results in increased IFN-γ-dependent lung bleeding during *N. brasiliensis* larvae migration. **a**, Experimental approach used for *N. brasiliensis* p.c. and i.v. infections. For the p.c. route, mice were infected with 500 L3s as described in Fig. 1a; for the i.v. route, mice were injected with 200 L3s in sterile PBS. **b**, Total lung RNA from naïve (white circles), p.c.-(black circles) and i.v.-infected (pink triangles) C57BL/6J mice was analysed with the Mouse Myeloid Innate Immunity V2 Panel NanoString panel and total sample variation was summarised via principal component analysis (PCA) across all genes. (*n* = 4 mice per group) **c**, Differentially expressed genes (DEGs) were visualised using a volcano plot, genes significantly upregulated in p.c. infection (red) and i.v. (blue) had a *p* value of <0.05. **d, e**, Overrepresented pathways determined via Ingenuity Pathway Analysis (IPA) in DEGs from p.c.-infected vs naïve controls **(d)** and i.v.-infected vs naïve controls **(e). f**, Flow cytometry analysis of intracellular IL-17A and IFN-γ in γδ T cells isolated from the lungs of naïve (white circles), p.c. (black circles) and i.v. (pink triangles) *N. brasiliensis*-infected C57BL/6J mice and stimulated ex vivo for 4h. Summary graph showing the percentages of IL-17A^+^ and IFN-γ^+^ cells within the γδ T cell population (*n* = 9-11 mice per group; pool of 2 independent experiments). **g**, Frequencies of Vγ6 cells (defined as Vγ1^-^Vγ4^-^) within the γδ T cell population isolated from the lungs of naïve (white circles), p.c. (black circles) and i.v. (pink triangles) *N. brasiliensis*-infected C57BL/6J mice (*n* = 8-10 mice per group; pool of 2 independent experiments). **h**, Frequencies of neutrophils within total leukocytes in the lungs of naïve (white circles), p.c. (black circles) and i.v. (pink triangles) *N. brasiliensis*-infected C57BL/6J mice (*n* = 10-14 mice per group; pool of 3 independent experiments). **i**, Optical density of the BAL of naïve (white circles), p.c. (black circles) and i.v. (pink triangles) *N. brasiliensis*-infected C57BL/6J mice measured at 400nm (*n* = 15-21 mice per group; pool of 4 independent experiments). **j**, *Ifngr* ^-/-^ and their WT counterparts were infected with 200 *N. brasiliensis* L3s i.v. **k**, Optical density of the BAL of naïve (white triangles), and *N. brasiliensis*-infected *Ifngr* ^+/+^ (black triangles) and *Ifngr* ^-/-^ (yellow triangles) mice measured at 400nm (*n* = 6 mice per group). **f-k**, Error bars represent mean±SD. Normality of the samples was assessed with Shapiro-Wilk normality test; statistical analysis was then performed using Kruskal-Wallis or one-way ANOVA test **(f, k)**. * *p* < 0.05; ** *p* < 0.01; *** *p* < 0.001; **** *p* < 0.0001

### *Activation of lung γδ17 T cells requires* N.brasiliensis *larvae migration through the skin*

Next, to better understand the initial immune response of lung γδ T cells to migrating N. brasiliensis larvae, at day 1 p.i. we sorted all the major lung populations of innate lymphocytes in p.c.- and i.v.-infected mice and performed single-cell RNA sequencing (scRNAseq) analysis **(Fig. 3a)**. We identified and annotated 16 distinct cell subsets across all treatment groups **(Fig. 3b)**. *N. brasiliensis* p.c. and i.v. infections led to discrete changes in the immune populations analysed when compared to naïve mice **(Fig. 3c)**. Relative frequencies of γδ T cell populations confirmed our flow cytometry data, with a marked decrease in naïve and γδ1 T cells in p.c.-infected mice, juxtaposed by an increase in γδ1 T cells observed during i.v. infection **(Fig. 3d)**. Moreover, across different populations of IFN-γ-producing cells (with the exception of ILC1), p.c. infection led to a decrease in the expression of a type-1 effector gene module **(Extended Data Table 1)**, when compared to their i.v.-infected counterparts and naïve controls **(Fig. 3e)**. Conversely, all IFN-γ-producing populations from i.v.-infected mice displayed an activation of the same type-1 effector gene module **(Fig. 3e)**. With regards to IL-17-producing cells, *N. brasiliensis* p.c. infection induced an increase in the relative frequencies of Vγ6^+^ and Vγ6^-^ γδ17 T cells **(Fig. 3d)**, whereas i.v. infections led to a specific decrease in Vγ6^+^ γδ17 T cells **(Fig. 3d)**. None of the γδ17 T cells from the lung expressed genes commonly associated with circulating T cells, such as *Sell* (encoding CD62L) and *Ccr7* (encoding CCR7; **Extended Data Fig. 3a**). This provides further evidence that the increase of γδ17 T cells within the lung tissue following *N. brasiliensis* p.c. infection are not recirculating cells. Furthermore, relative to naïve mice, Vγ6^+^ and Vγ6^-^ γδ17 T cells from i.v.-infected animals displayed only 3 and 1 differentially expressed genes (DEGs) respectively **(Fig. 3f, g)**, whereas p.c. infection led to 44 DEGs in Vγ6^+^ γδ17 T cells **(Fig. 3f, h)** and 29 DEGs in Vγ6^-^ γδ17 T cells **(Fig. 3g and Extended Data Fig. 3b)**. Notably, upregulated DEGs from p.c. Vγ6^+^ γδ17 T cells were associated with pathways related to T cell activation **(Fig. 3i)**; downregulated DEGs, on the other hand, were associated with positive regulation of cell–substrate adhesion **(Fig. 3j)**. Taken together, our results indicate that activation of γδ17 T cells in the lungs during *N. brasiliensis* infection only happens when the larvae migrate through the skin, which also induces the suppression of IFN-γ-producing cells. Bypassing the skin phase of the infection leads to an enrichment in lung type-1 cells and an increase in IFN-γ-biased transcriptional signatures.

**Fig. 3.**
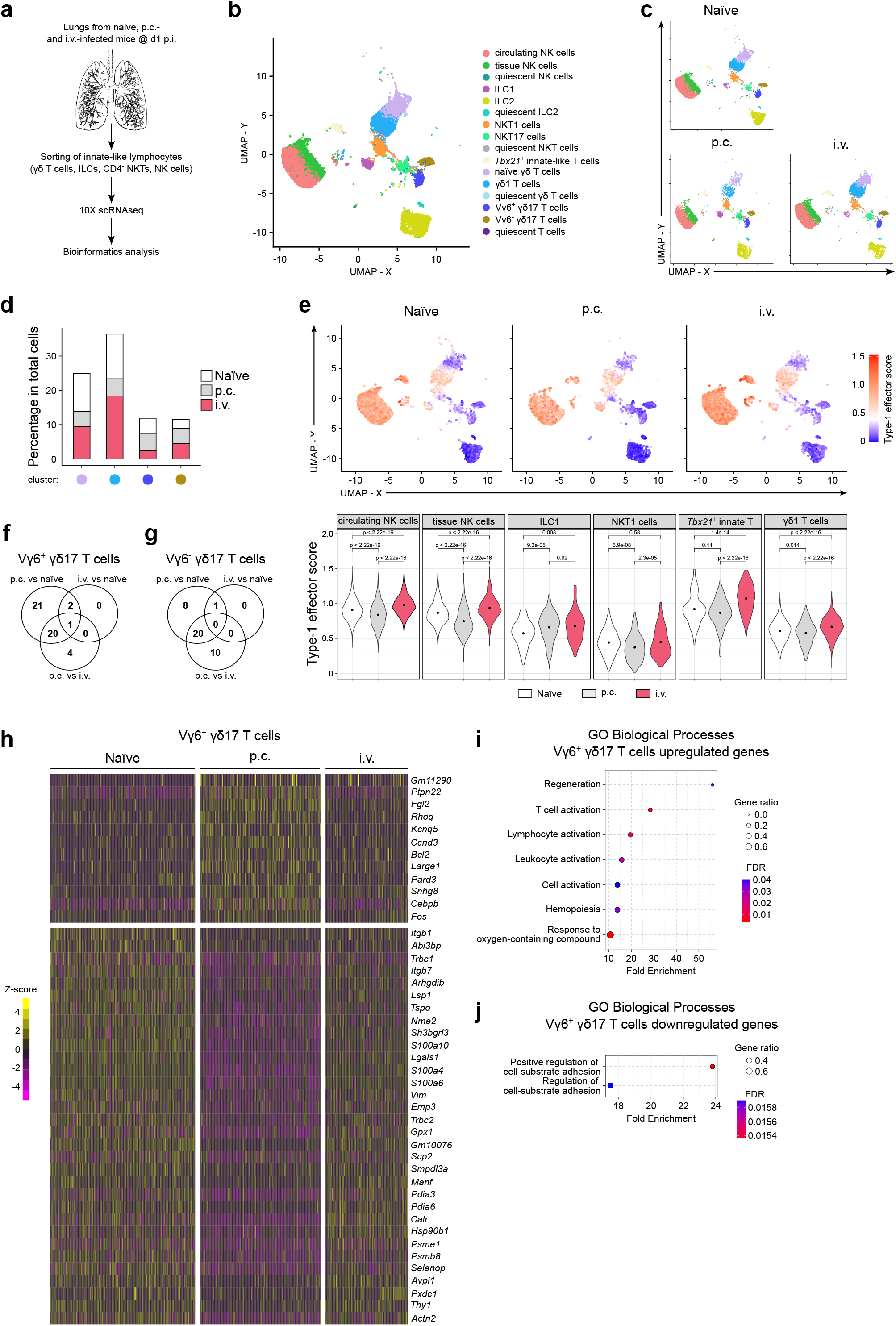
P.c., but not i.v., *N. brasiliensis* infection induces the activation of Vγ6^+^ γδ17 T cells and suppression of IFN-γ-producing innate lymphocytes. **a**, Schematic showing the strategy used for sorting innate lymphocytes prior to scRNAseq. **b**,**c**, Manually annotated Louvain clusters projected onto two dimensional UMAP plot **(b)** and separated by infection status **(c). d**, Percentage of total cell barcodes within different infection groups for naïve γδ T cells (lilac circle), γδ1 T cells (light blue circle), Vγ6^+^ γδ17 T cells (dark blue circle), Vγ6^-^ γδ17 T cells (golden circle) clusters. **e**, Type-1 effector signature was generated from a list of manually curated genes available from the literature(27–29) and calculated average standard scores (z-score) of this gene module were used to colour cells projected on a UMAP plot (red = high, blue = low). Summary violin plots showing the per-cell Type-1 effector score across different cell clusters and treatment conditions (Naïve = white; p.c. = grey; i.v. = pink). **f, g**, Venn diagram showing the overlap of DEGs (defined as adjusted *p* value <0.05, absolute log2 fold change of >1 and mean log2 expression value >1) between different infection route comparisons for Vγ6^+^ **(f)** and Vγ6^-^ γδ17 T cells **(g). h**, Heatmap showing the DEGs in Vγ6^+^ γδ17 T cells from naïve, p.c.- and i.v.-infected animals. Scale represents scaled and centred z-scores. **i**,**j**, Overrepresented GO: biological processes in upregulated **(i)** and downregulated **(j)** genes in Vγ6^+^ γδ17 T cells from p.c.-infected animals. **b-j**, scRNAseq data consists of sorted cells pooled from 4 mice per group.

### N. brasiliensis *induces the activation of migratory pathways in dermal* γδ *T cells*

As some of the enriched pathways in the Vγ6^+^ γδ17 T cells from the lungs of p.c.-infected mice suggested decreased ECM adhesion, and thus increased motility, we hypothesised that these cells were being activated elsewhere. We therefore assessed the effect of invading larvae on the cellular immune responses taking place in the skin, in particular by γδ T cells **(Fig. 4a)**. In notable contrast to what we observed in the lungs, intracellular staining of skin cells stimulated ex vivo showed no differences in IL-17A^+^ and IL-17F^+^ dermal γδ T cells between naïve and *N. brasiliensis*-infected mice **(Fig. 4b)**. Likewise, at 6h p.i. *N. brasiliensis* infection did not induce an increase of IL-17A or IL-17F in skin tissue lysates **(Extended Data Fig. 4a)**. Instead, *N. brasiliensis* larvae induced an increase in the skin levels of IL-6 and type-2 cytokines such as TSLP, IL-33, IL-5 and Ym1 **(Extended Data Fig. 4a, b)**. Despite the lack of differences in the production of IL-17 cytokines, infected mice displayed an increase in neutrophils (but not in monocytes) in the skin already at 6h p.i., being found mainly around *N. brasiliensis* larvae **(Extended Data Fig. 4c)**. At 24h p.i., infected mice showed marked changes in the immune cell populations present in the skin **(Fig. 4c and Extended Data Fig. 4d)**, with monocytes and neutrophils being the only two populations increased upon infection **(Fig. 4c and Extended Data Fig. 4d)**, as previously described(30). Interestingly, dermal γδ T cells were found to be decreased in infected mice **(Fig. 4c)**, further indicating a possible relocation of these cells.

**Fig. 4.**
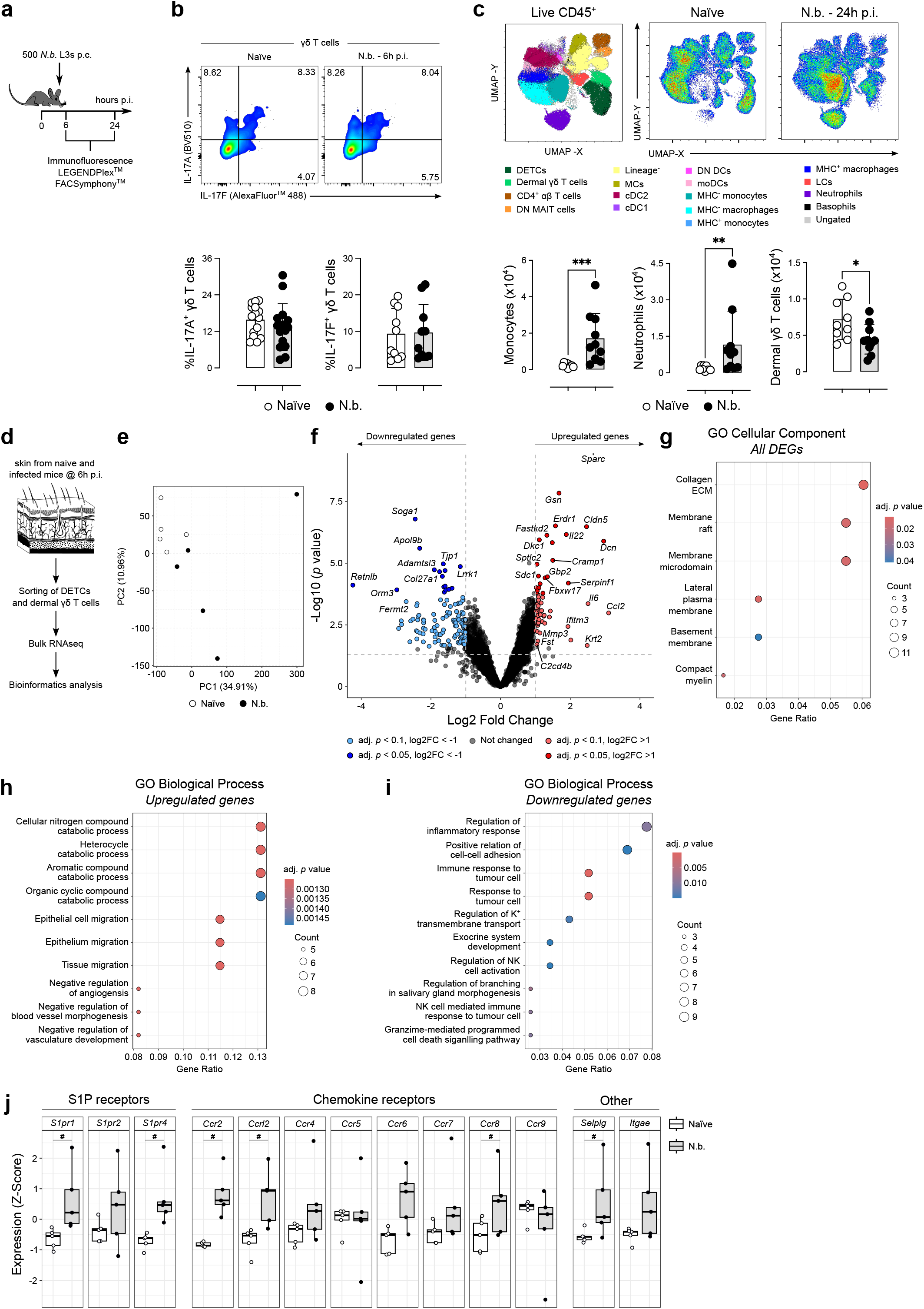
*. N. brasiliensis* larvae induce a migratory transcriptional program in dermal γδ T cells. **a**, C57BL/6J mice were infected with 500 *N. brasiliensis* L3s p.c. and analysed at 6h and 24h p.i. **b**, Flow cytometry analysis of intracellular IL-17A and IL-17F in dermal γδ T cells isolated from the skin of naïve (white circles) and *N. brasiliensis*-infected (black circles) mice and stimulated ex vivo for 3h. Summary graph showing the percentages of IL-17A^+^ and IL-17F^+^ cells within the γδ T cell population (*n* = 10-15 mice per group; pool of 3 independent experiments for IL-17A and pool of 2 independent experiments for IL-17F). **c**, Representative UMAP plot (pool of 5 mice per group) summarising the flow cytometry analysis of skin immune cell populations (left) and density UMAP plots summarising the data for naïve (middle) and *N. brasiliensis*-infected (right) mice. Summary graph showing the numbers of skin monocytes, neutrophils and dermal γδ T cells (*n* = 9-10 mice per group; pool of 2 independent experiments). **d**, Experimental approach employed to obtain the transcriptional signature of DETCs and dermal γδ T cells from naïve and *N. brasiliensis*-infected C57BL/6J mice. (*n* = 5 mice per group) **e**, Total sample variation in dermal γδ T cells from naïve (white circles) and *N. brasiliensis*-infected (black circles) mice was summarised via PCA across all genes. **f**, DEGs from dermal γδ T cells were visualised using a volcano plot, genes significantly upregulated (red), and downregulated (blue) during *N. brasiliensis* infection were defined as having an adjusted *p* value of <0.05 or <0.1 and absolute log2 fold change of >1. **g**, Overrepresented gene ontology (GO): cellular components in DEGs from dermal γδ T cells from *N. brasiliensis*-infected (black circles) mice. **h**, Overrepresented GO: biological processes in upregulated genes in dermal γδ T cells from *N. brasiliensis*-infected (black circles) mice. **i**, overrepresented GO: biological processes in downregulated genes in dermal γδ T cells from *N. brasiliensis*-infected (black circles) mice. **j**, normalised counts for specific genes converted to Z-scores. **b**,**c**, Error bars represent mean±SD. Normality of the samples was assessed with Shapiro-Wilk normality test accordingly; statistical analysis was then performed using Student’s t test or Mann-Whitney test. * *p* < 0.05; ** *p* < 0.01; *** *p* < 0.001; adjusted *p* value < 0.1

In order to investigate the effects of *N. brasiliensis* larvae on skin γδ T cells, we sorted both dendritic epidermal T cells (DETCs) and dermal γδ T cells from naïve and *N. brasiliensis*-infected mice at 6h p.i. for subsequent RNA sequencing **(Fig. 4d)**. Even within the short timeframe selected, *N. brasiliensis* infection induced significant changes in the transcriptional profile of dermal γδ T cells **(Fig. 4e)**, with almost 200 differentially expressed genes (DEGs) between naïve and infected mice **(Fig. 4f and Extended Data Table 2)**. Notably, most canonical γδ17 T cell activation genes, such as *Rorc, Il17a, Il23r, Gpr183* and *Blk*, were not significantly different between dermal γδ T cells from naïve and infected mice **(Fig. 4f and Extended Data Table 2)**, corroborating our protein data **(Fig. 4b and Extended Data Fig. 4a)** and suggesting the involvement of other signalling pathways. In fact, DEGs from dermal γδ T cells in *N. brasiliensis*-infected mice were found to be involved mostly with cell membrane and ECM components **(Fig. 4g)**. Accordingly, we found pathways related to cell migration (regulated by genes such as *Sparc, Dcn* and *Cd63*) and catabolic processes (mediated by genes such as *Il6, Fastkd2* and *Dkc1*) to be enriched in upregulated DEGs **(Fig. 4h)**. Downregulated DEGs were found to be mainly associated with the regulation of cytotoxic pathways (e.g., *Nkg7, Prf1* and *Havcr2*) and cell-cell adhesion (e.g., *Tjp1, Cdh1* and *Cd160*; **Fig. 4i**). Furthermore, dermal γδ T cells from infected mice showed increased gene expression of the S1P receptors *S1pr1* and *S1pr4*, and the chemokine receptor *Ccr2*, all of which have been shown to regulate γδ17 T cell migration during inflammation(31, 32) **(Fig. 4j)**. Hence, upon *N. brasiliensis* infection dermal γδ T cells engage a transcriptional programme directed to cell migration. Importantly, these changes seemed to be restricted to dermal γδ T cells, as DETCs from *N. brasiliensis*-infected mice showed no relevant transcriptional differences when compared to their naïve counterparts **(Extended Data Fig. 4e)**. Collectively, our analysis of the early immune events in the skin of *N. brasiliensis*-infected mice suggests dermal γδ T cells become activated and engage a migratory programme, leading to their disappearance from the skin after 24h p.i.

### *Dermal γδ T cells migrate to the lungs early during* N. brasiliensis *infection*

Our data implied that the increase in γδ17 T cells in the lungs during *N. brasiliensis* infection could stem from the migration of dermal γδ T cells and not local expansion – as Ki67 staining showed no differences in the proliferation status of lung γδ17 T cells between naïve and *N. brasiliensis*-infected mice at day 1 p.i. **(Extended Data Fig. 5a)**. In order to track migratory γδ T cells, we initially employed photoconvertible Kaede mice. However, we were unable to observe an immune response to invading *N. brasiliensis* lar-vae using Kaede mice, as evidenced by the lack of changes in skin draining lymph node cell numbers and no neutrophil recruitment to the lungs **(Extended Data Fig. 5b)**. Therefore, our skin infection model was incompatible with this mouse strain likely due to the immunosuppressive effects of UV radiation on the skin(33). As an alternative, we employed an inducible *Id2*^CreERT2^x*Rosa26*^RFP^ reporter mouse line, coupled with topical administration of 4-hydroxytamoxifen (4OH-T) on the skin area where the *N. brasiliensis* infection would later take place **(Fig. 5a)**. As Cre recombinase activation in distal tissues upon topical 4OH-T application is minimal(34), this strategy allows for local cell-specific genetic manipulation. We achieved a sizeable proportion of innate-like lymphocytes – including dermal γδ T cells – expressing the red fluorescent protein RFP **(Fig. 5b and Extended Data Fig. 5c)**. Using this approach, we observed the expected decrease in dermal γδ T cell numbers in the skin **(Fig. 5c)** and increase in IL-17A^+^ IL-17F^-^ and IL-17A^+^IL-17F^+^ γδ T cells in the lungs upon *N. brasiliensis* infection **(Fig. 5d)**. To overcome the variability of the recombination process, we normalised the RFP percentages in distal organs to the penetrance of Cre recombinase in the skin of each mouse. Thus, we observed an increase frequency of RFP expression within total lung γδ T cells **(Fig. 5e)**, together with an increase in the numbers of RFP^+^ γδ T cells **(Fig. 5f)**. This increase was mostly driven by the accumulation of RFP^+^ cells within the population of Vγ6, and not Vγ4^+^ or Vγ1^+^, γδ T cells **(Fig. 5g)**. Critically, we found an increased proportion and total numbers of IL-17A^+^RFP^+^ cells within γδ T cells from *N. brasiliensis*-infected mice when compared to naïve controls **(Fig. 5h and 5i)**, corroborating our hypothesis that, in this model, activated γδ17 T cells in the lungs originate from the skin. Of note, *N. brasiliensis* infection did not induce an increase in RFP^+^ γδ T cells in the ear draining lymph nodes at the timepoint analysed **(Extended Data Fig. 5d)**. As an further approach, we employed *Trdc*^CreERT2^x*Rosa26*^RFP^ reporter mice using a similar protocol as described above **(Extended Data Fig. 5e)** to specifically track skin γδ T cells. Despite lower Cre recombinase penetrance in these mice **(Extended Data Fig. 5f)**, we observed a smaller, but significant, increase in the proportion of RFP^+^ γδ T cells in the lungs of *N. brasiliensis*-infected mice compared to controls **(Extended Data Fig. 5g)**.

**Fig. 5.**
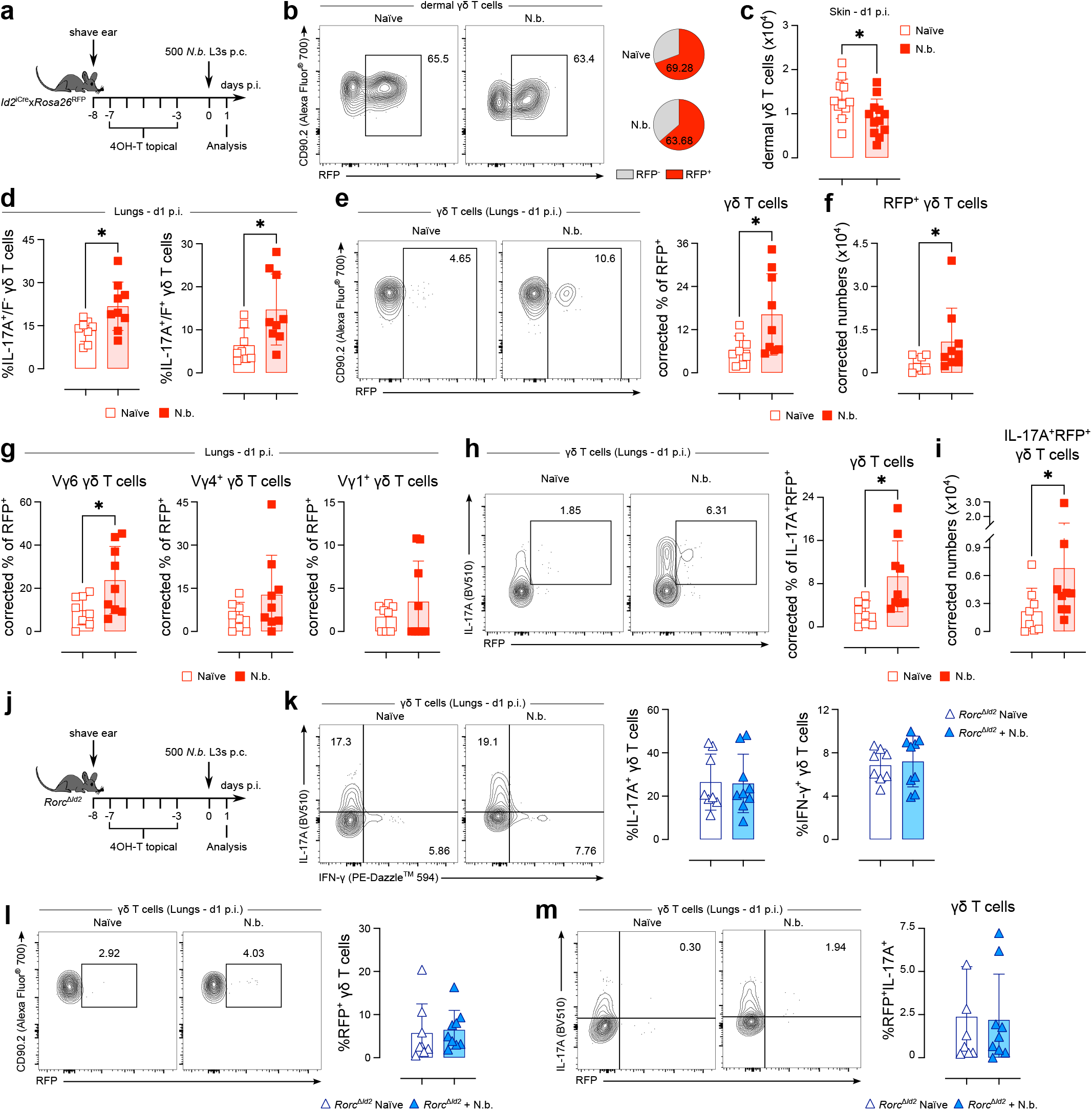
Migratory dermal γδ17 T cells are responsible for the expansion in the lung γδ17 T cell population during *N. brasiliensis* infection. **a**, *Id2*^CreERT2^x*Rosa26*^RFP^ mice had their right ear shaved, and starting on the following day were treated for five consecutive days with topical 4OH-T. Mice were then rested for 3 days before being infected with 500 *N. brasiliensis* L3s p.c. **b**, Flow cytometry analysis of RFP expression in dermal γδ T cells isolated from the skin of naïve and *N. brasiliensis*-infected mice. Summary pie charts showing the percentage of RFP^+^ cells (red slice) within dermal γδ T cells (*n* = 11-13 mice per group; pool of 3 independent experiments). **c**, Summary graph showing the numbers of dermal γδ T cells isolated from the skin of naïve (white squares) and *N. brasiliensis*-infected (red squares) mice (*n* = 11-12 mice per group; pool of 3 independent experiments). **d**, Summary graph showing the percentages of IL-17A^+^IL-17F^-^ and IL-17A^+^IL-17F^+^ cells within the γδ T cell population in the lungs of naïve (white squares) and *N. brasiliensis*-infected (red squares) mice (*n* = 8-9 mice per group; pool of 3 independent experiments). **e, f**, Flow cytometry analysis of RFP expression in lung γδ T cells showing the corrected percentage **(e)** and numbers **(f)** of RFP^+^ γδ T cells isolated from the lungs of naïve (white squares) and *N. brasiliensis*-infected (red squares) *Id2*^CreERT2^x*Rosa26*^RFP^ mice (*n* = 8-9 mice per group; pool of 3 independent experiments). **g**, Summary graphs showing the corrected percentages of RFP^+^ cells within Vγ6 (defined as Vγ1^-^Vγ4^-^), Vγ1^-^Vγ4^+^ and Vγ1^+^Vγ4^-^ γδ T cells isolated from the lungs of naïve (white squares) and *N. brasiliensis*-infected (red squares) *Id2*^CreERT2^x*Rosa26*^RFP^ mice (*n* = 8-9 mice per group; pool of 3 independent experiments). **h, i**, Flow cytometry analysis of RFP and IL-17A coexpression in lung γδ T cells showing the corrected percentages **(h)** and numbers **(i)** of IL-17A^+^RFP^+^ γδ T cells isolated from the lungs of naïve (white squares) and *N. brasiliensis*-infected (red squares) *Id2*^CreERT2^x*Rosa26*^RFP^ mice (*n* = 8-9 mice per group; pool of 3 independent experiments). **j**, *Id2*^CreERT2^x*Rorc*^fl/fl^ mice had their right ear shaved, and starting on the following day were treated for five consecutive days with topical 4OH-T. Mice were then rested for 3 days before being infected with 500 *N. brasiliensis* L3s p.c. **k**, Flow cytometry analysis of intracellular IL-17A and IFN-γ in γδ T cells isolated from the lungs of naïve (white triangles), and *N. brasiliensis*-infected (blue triangles) *Id2*^CreERT2^x*Rorc*^fl/fl^ mice and stimulated ex vivo for 4h. Summary graph showing the percentages of IL-17A^+^ and IFN-γ^+^ cells within the γδ T cell population (*n* = 6-9 mice per group; pool of 2 independent experiments). **l, m**, Flow cytometry analysis of RFP expression and summary graphs showing the percentage of RFP^+^ cells within total γδ T cells **(l)** and IL-17A^+^RFP^+^ γδ T cells **(m)** isolated from the lungs of naïve (white triangles) and *N. brasiliensis*-infected (blue triangles) *Id2*^CreERT2^x*Rorc*^fl/fl^ mice (*n* = 6-9 mice per group; pool of 2 independent experiments). **e-i**, Individual percentages of RFP^+^ cells were normalised to the respective Cre penetrance calculated from skin percentages of RFP^+^ cells, assuming 100% recombination efficiency. **c-i**, Error bars represent mean±SD. Normality of the samples was assessed with Shapiro-Wilk normality test; statistical analysis was then performed using Student’s t test or Mann-Whitney test. * *p* < 0.05.

To further confirm a role for dermal γδ17 T cells in the establishment of lung type-17 immune responses early during *N. brasiliensis* larvae migration, we used *Id2*^CreERT2^ mice crossed with *Rorc*^fl^ animals to deplete IL-17-producing innate lymphocytes (*Rorc*^Δ*Id2*^; **Fig. 5j and Extended Data Fig. 5h**). Upon N. brasiliensis infection, *Rorc*

To gain insight into the mechanisms underlying dermal γδ17 T cell migration, we prevented cell trafficking by employing transgenic mouse models and small molecule inhibitors. γδ T cell migration has been shown to be dependent on S1PR(31), and we observed an increased expression of *S1pr1* and *S1pr4* in dermal γδ T cells upon *N. brasiliensis* infection **(Fig. 4j)**. Therefore, we blocked leukocyte trafficking by using the S1PR modulator FTY720 in our experimental model **(Fig. 6a)**. In contrast to untreated animals, FTY720-treated mice did not display either the expected increase in lung Vγ6 γδ17 T cells **(Fig. 6b, c and Extended Data Fig. 6a)** nor did they exhibit a significant expansion of the Vγ6 population upon *N. brasiliensis* infection **(Fig. 6d and Extended Data Fig. 6b, c)**. Accordingly, FTY720-treated mice failed to recruit neutrophils to the lungs upon *N. brasiliensis* infection, in a clear contrast to their untreated counterparts **(Fig. 6e)**. This notwithstanding, FTY720-treatment was not sufficient to alter lung haemorrhage **(Fig. 6f)**. CCR2 has also been shown to mediate γδ T cell migration during inflammation(32), and we observed an increase in *Ccr2* gene expression in dermal γδ T cells upon *N. brasiliensis* infection **(Fig 4j)**. While as expected, naïve *Ccr2*^-/-^ animals were deficient in lung monocytes and interstitial macrophages when compared to their WT counterparts **(Extended Data Fig. 6d, e)**, T cells, and specifically γδ T cell subpopulations, were not affected by CCR2 deficiency, indicating that in the time-point analysed this model was well-suited to investigate the role of CCR2 in γδ T cell migration **(Extended Data Fig. 6d-f)**. Thus, we infected *Ccr2*^-/-^ mice and their WT littermates with *N. brasiliensis* and analysed their skin and lung immune responses **(Fig. 6g)**. Whereas infected WT mice displayed the expected decrease in dermal γδ T cells in the skin upon *N. brasiliensis* infection, their CCR2-deficient counterparts failed to do so **(Fig. 6h)**. Likewise, infected WT mice displayed an increase in γδ17 T cells in the lungs together with a decrease in IFN-γ-producing γδ T cells **(Fig. 6i)**, in contrast to what was observed for *Ccr2*^-/-^ mice **(Fig. 6j)**. *N. brasiliensis* infection also induced a shift in Vγ usage, with a marked increase in Vγ6 cells within WT, but not in *Ccr2*^-/-^, lung γδ T cells **(Fig. 6k and Extended Data Fig. 6g, h)**. Critically, whereas *N. brasiliensis*-infected *Ccr2*^+/+^ mice displayed the expected increase in lung neutrophils, this was not observed in *Ccr2*^-/-^ animals **(Fig. 6l)**. Nonetheless, *Ccr2*^-/-^ mice ex-hibited more BAL haemorrhage than their WT counterparts **(Fig. 6m)**. Altogether, our results reveal that dermal Vγ6 γδ17 T cells migrate from the skin to the lungs, in a CCR2- and S1PR-dependent manner, early during *N. brasiliensis* infection to kickstart IL-17-mediated immune responses with impacts on parasite-induced lung damage.

**Fig. 6.**
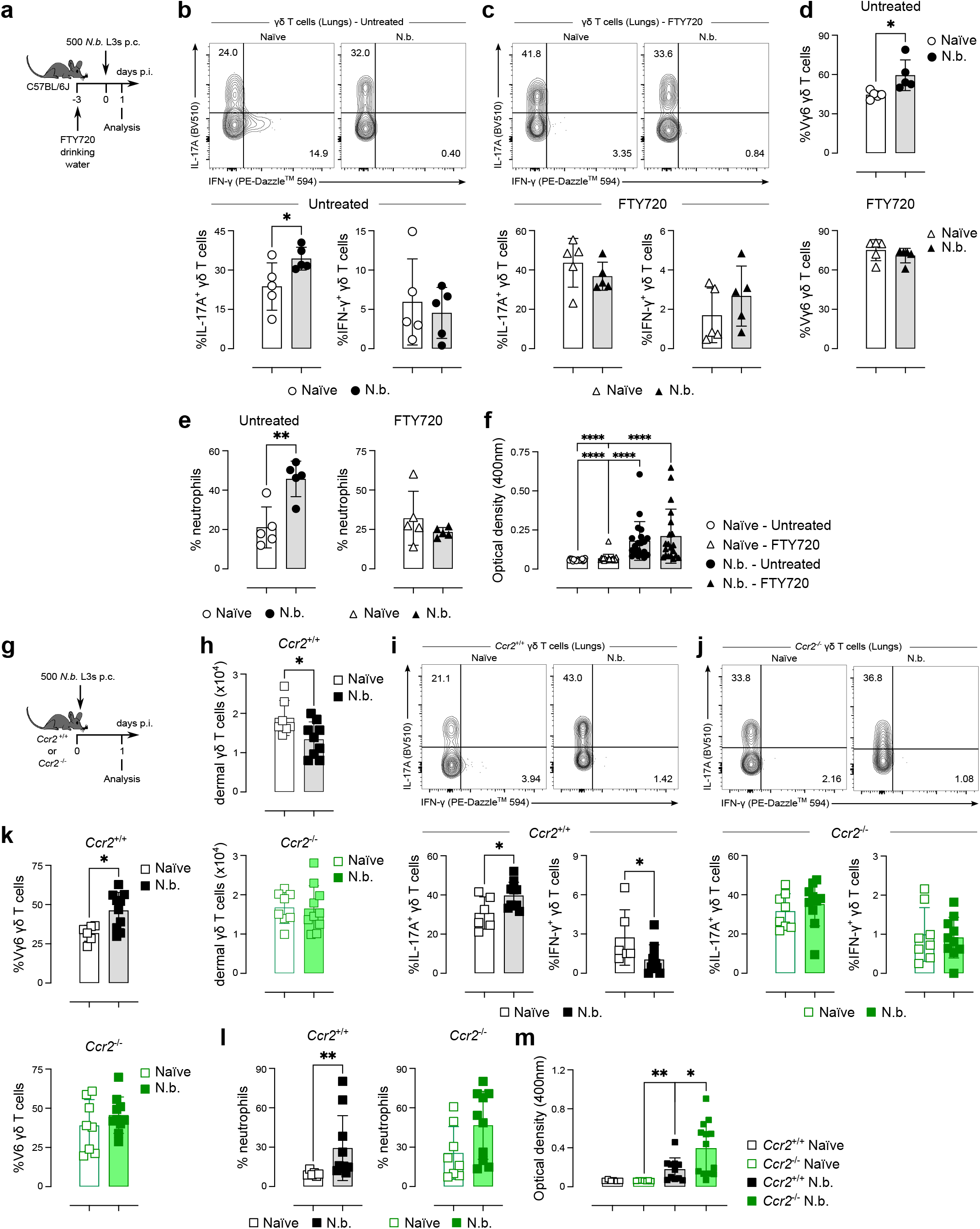
Dermal γδ17 T cells migrate to the lungs during *N. brasiliensis* infection in a S1PR- and CCR2-dependent fashion. **a**, C57BL/6J mice were treated with FTY720 in the drinking water starting 3 days prior to the infection, and were then infected or not with 500 *N. brasiliensis* L3s p.c.; mice were analysed at day 1 p.i. Control mice were kept with untreated drinking water **b, c**, Summary graphs showing the percentages of IL-17A^+^ and IFN-γ^+^ cells within lung γδ T cells from untreated **(b)** and FTY720-treated **(c)** naïve (white symbol) and *N. brasiliensis*-infected (black symbol) mice after 4h of ex vivo stimulation (*n* = 5 mice per group; representative of at least 3 independent experiments). **d**, Frequencies of Vγ6 cells (defined as Vγ1^-^Vγ4^-^) within the γδ T cell population isolated from the lungs of untreated (top) and FTY720-treated (bottom) naïve (white symbols) or *N. brasiliensis*-infected C57BL/6J mice (black symbols) (*n* = 5 mice per group; representative of at least 3 independent experiments). **e**, Frequencies of neutrophils within total leukocytes in the lungs of untreated (left) and FTY720-treated (right) naïve (white symbols) or *N. brasiliensis*-infected C57BL/6J mice (black symbols) (*n* = 5 mice per group; representative of at least 3 independent experiments). **f**, Optical density of the BAL of untreated and FTY720-treated naïve (white symbols) or *N. brasiliensis*-infected C57BL/6J mice (black symbols) (*n* = 19-20 mice per group; pool of at 4 independent experiments). **g**, *Ccr2*^-/-^ and *Ccr2*^+/+^ mice were infected 500 *N. brasiliensis* L3s p.c. and were analysed at day 1 p.i. **h**, Summary graph showing the numbers of dermal γδ T cells isolated from the skin of naïve (open squares) and *N. brasiliensis*-infected (filled squares) *Ccr2*^+/+^ (top) and *Ccr2*^-/-^ (bottom) mice (*n* = 7-10 mice per group; pool of 3 independent experiments). **i, j**, Flow cytometry analysis of intracellular IL-17A and IFN-γ in γδ T cells isolated from the lungs of naïve (open squares) and *N. brasiliensis*-infected (filled squares) *Ccr2*^+/+^ **(i)** and *Ccr2*^-/-^ **(j)** mice and stimulated ex vivo for 4h. Summary graph showing the percentages of IL-17A^+^ and IFN-γ^+^ cells within γδ T cells (*n* = 6-10 mice per group; pool of 3 independent experiments). **k**, Summary graph showing the percentages Vγ6 cells (defined as Vγ1^-^Vγ4^-^) within γδ T cells isolated from the lungs of naïve (open squares) and *N. brasiliensis*-infected (filled squares) *Ccr2*^+/+^ (top) and *Ccr2*^-/-^ (bottom) mice (*n* = 6-10 mice per group; pool of 3 independent experiments). **l**, Frequencies of neutrophils within total leukocytes in the lungs of naïve (open squares) and *N. brasiliensis*-infected (filled squares) *Ccr2*^+/+^ (left) and *Ccr2*^-/-^ (right) mice (*n* = 6-10 mice per group; pool of 3 independent experiments). **m**, Optical density of the BAL of naïve (open squares) and *N. brasiliensis*-infected (filled squares) *Ccr2*^+/+^ (left) and *Ccr2*^-/-^ (right) mice (*n* = 6-13 mice per group; pool of 3 independent experiments). Error bars represent mean±SD. Normality of the samples was assessed with Shapiro-Wilk normality test; statistical analysis was then performed using Student’s t test or Mann-Whitney test **(b-e, h-k)**, or Kruskal-Wallis or one-way ANOVA test **(f, m)**. * *p* < 0.05; ** *p* < 0.01; **** *p* < 0.0001

### *IL-1R signalling underlies dermal γδ T cell activation and migration to the lungs during* N. brasiliensis *infection*

Having shown that the expansion in γδ17 T cells observed in the lungs during the early phase of N. brasiliensis infection stemmed from migratory dermal Vγ6 γδ T cells, we next sought to understand the signals driving their activation in the skin. Similar to other unconventional T lymphocytes, the role of TCR signalling in the activation of γδ T cells is still a matter of debate(15). To examine if *N. brasiliensis* larvae were inducing TCR activation in dermal Vγ6 γδ T cells we employed the Nur77-Tempo mouse line that allows temporal identification of cells that have engaged TCR signalling(35). Recent TCR activation (<4 to 7 hours) leads to the expression of the unstable form of the Fluorescent Timer (FT) protein (FT Blue), whereas historic TCR signalling is characterised by the stable form of the FT protein **(FT Red; Extended Data Fig. 7a)**(35). Total dermal γδ T cells from *N. brasiliensis*-infected mice did not show any sign of TCR engagement at 4h p.i. **(Fig. 7a)**. Likewise, both the Vγ6 and Vγ4^+^ subsets of dermal γδ T cells failed to display any indication of TCR activation upon infection **(Extended Data Fig. 7b, c)**.

**Fig. 7.**
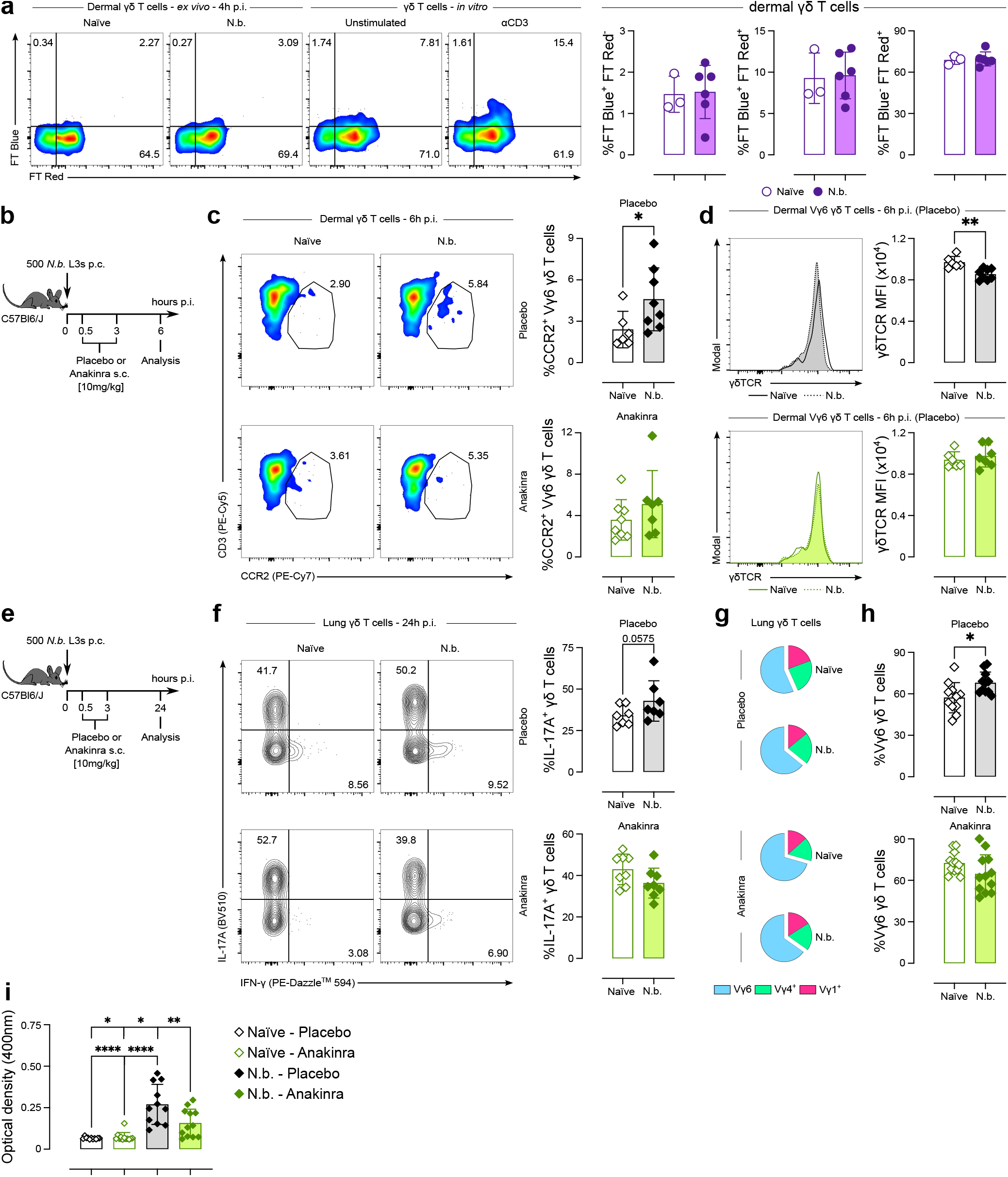
Dermal γδ T cells are activated via the IL-1R signalling pathway during *N. brasiliensis* infection. **a**, Nur77-Tempo mice were infected or not with 500 *N. brasiliensis* L3s p.c. and analysed at 4h p.i. Flow cytometry analysis and summary graphs showing FT protein expression among dermal γδ T cells from naïve (white circles) and *N. brasiliensis* infected (purple circles) mice. As a positive control of FT Blue expression lymph node cells were stimulated in vitro with 2.5 µg/mL of anti-CD3 for 4h and FT Blue expression was analysed on γδ T cells (*n* = 3-6 mice per group; representative of 2 independent experiments). **b**, C57BL/6J mice were infected with 500 *N. brasiliensis* L3s p.c. and treated with 100 mg/g of Anakinra or placebo at 30 minutes p.i.; a second dose was given at 3h p.i.; mice were then analysed at 6h p.i. **c**, Flow cytometry analysis and summary graphs showing CCR2 protein expression among dermal Vγ6 γδ T cells (defined as Vγ1^-^Vγ4^-^) from naïve (white diamonds) and *N. brasiliensis* infected (filled diamonds) mice treated with placebo (top) or Anakinra (bottom) (*n* = 6-8 mice per group; pool of 2 independent experiments). **d**, Flow cytometry analysis and summary graphs showing the median fluorescence intensity of surface γδTCR among dermal Vγ6 γδ T cells (defined as Vγ1^-^Vγ4^-^) from naïve (full line in the histogram plots and white diamonds in the bar plots) and *N. brasiliensis* infected (dashed line in the histogram plots and filled diamonds in the bar plots) mice treated with placebo (top) or Anakinra (bottom) (*n* = 6-8 mice per group; pool of 2 independent experiments). **e**, C57BL/6J mice were infected with 500 *N. brasiliensis* L3s p.c. and treated with 100 mg/g of Anakinra or placebo at 30 minutes p.i.; a second dose was given at 3h p.i. in the first experiment, but not in the subsequent two; mice were then analysed at 24h p.i. **f**, Flow cytometry analysis of intracellular IL-17A and IFN-γ in γδ T cells isolated from the lungs of naïve (white diamonds) and *N. brasiliensis* infected (filled diamonds) mice treated with placebo (top) or Anakinra (bottom) and stimulated ex vivo for 4h. Summary graphs showing the percentages of IL-17A^+^ and IFN-γ^+^ cells within γδ T cells (*n* = 7-8 mice per group; pool of 2 independent experiments). **g**, Pie charts depicting the Vγ chain usage among lung γδ T cells from naïve (white diamonds) and *N. brasiliensis* infected (filled diamonds) mice treated with placebo (top) or Anakinra (bottom) (*n* = 11-12 mice per group; pool of 3 independent experiments). **h**, Summary graph showing the percentages Vγ6 cells (defined as Vγ1^-^Vγ4^-^) within γδ T cells isolated from the lungs of naïve (white diamonds) and *N. brasiliensis* infected (filled diamonds) mice treated with placebo (top) or Anakinra (bottom) (*n* = 11-12 mice per group; pool of 3 independent experiments). **i**, Optical density of the BAL of naïve (white diamonds) and *N. brasiliensis* infected (filled diamonds) mice treated with placebo (top) or Anakinra (bottom) (*n* = 11-12 mice per group; pool of 3 independent experiments). Error bars represent mean±SD. Normality of the samples was assessed with Shapiro-Wilk normality test; statistical analysis was then performed using Student’s t test or Mann-Whitney test (**a-h)**; or one-way ANOVA test **(i)**. * *p* < 0.05; ** *p* < 0.01; **** *p* < 0.0001

γδ17 T cells are known to be activated by the cytokines IL-1β and IL-23(15), with IL-1β being more involved in innate-like responses when compared to IL-23(36, 37). Therefore, to test if IL-1R signalling was involved in the activation of dermal γδ T cells during *N. brasiliensis* infection, we treated mice either with the IL-1R antagonist Anakinra or with a placebo formulation **(Fig. 7b)**. Dermal Vγ6 γδ T cells from placebo-treated mice displayed a significant increase in the expression of CCR2 (but not S1PR1) upon infection while treatment with Anakinra prevented the increase in CCR2 expression **(Fig. 7c and Extended Data Fig. 7d)**. Similarly, *N. brasiliensis* infection induced the decrease of γδTCR expression (a well-known characteristic of activated T cells) in dermal Vγ6 γδ T cells from the placebo-, but not the Anakinratreated group **(Fig. 7d)**. Apart from a trend for increased CCR2 expression in the placebo group, *N. brasiliensis* infection did not seem to induce the activation of Vγ4^+^ γδ T cells in the time point analysed **(Extended Data Fig. 7e-g)**. Analysis of the lungs of placebo- and Anakinra-treated mice at day 1 p.i. **(Fig. 7e)** showed the expected increase in γδ17 T cells and Vγ6 γδ T cells in control animals **(Fig. 7f-h and Extended Data Fig. 7h)**. Critically, IL-1R signalling blockade prevented the expansion of lung γδ17 T cells and Vγ6 γδ T cells **(Fig. 7f-h and Extended Data Fig. 7h)**. Finally, Anakinra treatment led to decreased BAL haemorrhage in response to lung migrating *N. brasiliensis* larvae **(Fig. 7i and Extended Data Fig. 7i)**. Thus, in summary, dermal Vγ6 γδ T cells are activated via the IL-1R signalling pathway during *N. brasiliensis* infection in a process seemingly independent of TCR activation. IL-1R blockade prevents the accumulation of γδ17 T cells in the lungs, ultimately affecting the tissue damage response.

## Discussion

Skin to lung migration represents an evolutionary common pathogen life cycle and thus tissue-migrating nematodes are valuable models for investigating the molecular cues underlying inter-tissue communication. Whereas pathogen type has traditionally been regarded as the driver of different types of immune responses, the tissue where the host-pathogen encounter occurs will also play a big part in this process(2). Here we show that during experimental *N. brasiliensis* infection, dermal γδ17 T cells are activated in the skin via IL-1R signalling and then migrate to the lungs, in a CCR2- and S1PR-dependent fashion, during the early stages of the parasite life cycle within the host. In the lung these cells orchestrate the initial events that shape the pulmonary immune response against *N. brasiliensis*: production of IL-17 and suppression of IFN-γ-producing cells. Bypassing the skin phase of infection, or blocking γδ T cell activation, precludes the migration of γδ17 T cells from the skin to the lungs, leading to marked changes in the pulmonary immune response to *N. brasiliensis*. Importantly, resident lung γδ17 T cells are not activated in loco by *N. brasiliensis* larvae arriving directly to the lungs; instead, i.v. infections induce the activation of IFN-γ-producing innate lymphocytes and increase tissue damage.

The induction of IL-17 production by lung γδ T cells during *N. brasiliensis* infection has always been a counterintuitive finding, especially when taking into consideration that chitin, a component of the nematode cuticle has the ability to activate lung ILC2s, which in turn suppress γδ17 T cells(38). Using different routes to establish *N. brasiliensis* infections, we found that the primary immune response to invading larvae is similar in both the skin and lungs, when they are the first point of contact between host and pathogen. Skin and lung possess obvious structural and functional differences in addition to discrete resident immune cell populations, arguing that pathogen-derived pathogen-associated molecular patterns (PAMPs) override tissue particularities in setting up the subsequent immune response. In particular, *N. brasiliensis* larvae seem to initially induce type-I and type-II IFN signalling together with type-2 immune responses on the body’s first encounter with larvae, whether skin or lung. This response, which would not normally take place in the lungs, leads to uncontrolled inflammation and increased lung bleeding. Of note, *N. brasiliensis* larvae ex-sheath after going through the skin(30), enhancing parasite fitness. In contrast, i.v. infection bypasses this step, with larvae likely ex-sheathing in the bloodstream or directly in the lungs, exposing the cuticle as the primary source of PAMPs. This shift in PAMP availability possibly contributes to the change in the initial immune response towards a type-1 / type-2 bias. Nonetheless, the migration of dermal γδ17 T cells to the lungs can be seen as a host mechanism to fine-tune the immune response against the migrating larvae, ultimately aimed at maintaining tissue integrity.

In the dermis, we find that γδ T cells are almost exclusively IL-17-producing cells(16). Although dermal γδ17 T cells are activated early after parasite invasion, *N. brasiliensis* larvae do not induce their expansion or an increase in IL-17 production, as was recently observed during *S. mansoni* infection(39, 40). Instead, we observed an increase in the expression of genes related to cell movement, such as chemokine receptors and cell – extracellular matrix interactions. Previously, we have shown that overexpression of Ym1 in the peritoneal cavity induces the activation of local γδ17 T cells via induction of IL-1β production(10). Here we show that IL-1R signalling is also key in mediating the activation, and consequent migration, of dermal γδ17 T cells. Interestingly, Ym1 was also induced in the skin upon *N. brasiliensis* larvae penetration; whether this molecule is upstream of IL-1β in this case is yet to be determined. IL-1R signalling has been shown to be essential for the activation of innate-like responses by dermal γδ17 T cells(36). Even though IL-23 is well known to act in synergy with IL-1β to induce IL-17 production by γδ T cells, its importance seems to be more marked in the activation of adaptive-like Vγ4^+^ γδT cells (which do not depend on IL-1R signalling)(37). The absence of TCR activation in dermal γδ T cells during *N. brasiliensis* infection reinforces the idea that these cells act in an innate-like fashion in this infection model. Dermal γδ T cells are known to be a motile population that exhibits a steady migration rate between the skin tissue and draining lymph nodes(17, 31). Upon inflammation, these cells can even reach distal skin sites(31, 41), but to our knowledge γδ T cell migration from the skin to other barrier sites has not yet been described. It is important to note that in humans chronically exposed to skin-penetrating hookworms, circulating γδ T cells display distinct functional signatures when compared to those from non-infected populations(42). Furthermore, γδ17 T cells from the gut have been shown to migrate to different organs(18, 43). Microbiota-activated small intestine γδ17 T cells migrate from the gut to the meninges during stroke and contribute to inflammation by recruiting neutrophils and monocytes(18). Similarly, small intestine memory γδ17 T cells were shown to migrate to the lungs during sepsis and contribute to acute lung injury through the pro-duction of IL-17A(43). Thus, it is conceivable that barrier-tissue-associated γδ17 T cells, particularly Vγ6^+^ γδ T cells, might support a mucosal-wide network.

An unexpected finding from this study was that IFN-γ, and not neutrophils, appears to play the greater role in driving lung haemorrhage. We and others have previously shown that depleting neutrophils, or preventing their recruitment ameliorates the alveolar lung injury induced by *N. brasiliensis*(11, 12, 24). The data presented here indicates that the enhanced bleeding primarily stems from physical damage caused by the migrating larvae with IFN-γ possibly further enhancing the haemorrhagic response. IFN-γ has previously been shown to increase vascular permeability both in vitro and in vivo(44, 45), and IFN-γ antibody blockade decreases acute lung injury in a murine flu infection model(26). Thus, our data suggests that upon encounter with the parasite in the skin dermal γδ17 T cells rapidly migrate to the lungs where they suppress, by yet unidentified mechanisms, type-1 effector subsets, mitigating acute haemorrhagic damage on parasite arrival. Nonetheless, IL-17 dependent neutrophilia does contribute to subsequent loss of alveolar structure12. Moreover, our data indicates that the complete absence of IL-17A and IL-17F (or blocking upstream activation signals) dampens both the immune response to *N. brasiliensis* larvae and acute lung damage. While seemingly at odds with the increased damage observed in i.v.-infected mice and *Ccr2*^-/-^ animals, in both these situations there are still IL-17-producing cells in the system. Therefore, our results point to a fine tuning of IL-17 levels (and possibly neutrophils) being the determinant between a subpar anthelminthic response and excessive tissue damage. This host adaptation might also be beneficial to (and maybe even induced by) the parasite, as ensuring host survival guarantees the maintenance of a productive parasitic life cycle. Accordingly, IL-17 deficiency has been shown to result in increased worm burden(10).

While the functional consequences of γδ17 T cell migration may vary with the context, our results show that the cellular make-up in non-lymphoid organs can be influenced by the immune events happening in distal sites. Our data using inducible fluorescent reporters further suggests that mi-gration of γδ T cells around mucosal sites occurs in steady state conditions, as even in the absence of an infection skin-derived cells could be found in the lungs. However, topical tamoxifen treatment alone might induce a low-grade inflammatory insult in the skin triggering the migration of dermal γδ17 T cells to the lungs albeit at a lower migration rate than that induced by *N. brasiliensis* infection. The atopic march theory postulates that airway allergies (often dependent on some form of type-2 immunity) arise from allergic responses taking place in the skin during early life(46). Thus, our results using an infectious trigger of type-2 immune responses could serve as a basis for identifying mechanisms behind the atopic march. In particular it would be interesting to determine if percutaneous allergen challenge promotes the migration of dermal γδ T cells, or other cell populations to the lungs. Overall, the unravelling of a skin – lung axis underlying inter-tissue communication during inflammatory responses opens exciting new avenues to understand immune responses at the whole-organismal level.

## Methods

### Mice and ethics

C57BL/6J^crl^ mice were obtained from Charles River. Id2^tm1.1(cre/ERT2)Blh^; B6(Cg)-Rorc^tm3Litt/J^; Gt(ROSA)26Sor^Ai14^ mice were originally provided by Dr. David Withers. Tcrd^tm1.1(cre/ERT2)Zhu^ were kindly provided by Dr. Adrian Hayday. Ccr2^tm1Ifc^ (*Ccr2*^-/-^) mice were kindly provided by Dr. Jo Konkel. *Ifngr*^-/-^ mice were kindly provided by Dr. Lizzie Mann(47). Kaede mice (Tg(CAG-tdKaede)15Utr) were kindly provided by Dr. Amy Saunders. Nur77-Tempo mice were generated and kindly gifted by Dr. David Bending(35). Male and female mice were age- and sex-matched and housed in individually ventilated cages. Experimental mice were not randomised in cages, but each cage was randomly assigned to a treatment group. Mice were culled by asphyxiation in a rising concentration of CO2. Experiments were performed in accordance with the United Kingdom Animals (Scientific Procedures) Act of 1986 (project license number 70/8547 and PP4115856).

### N. brasiliensis *infection*

*N. brasiliensis* was maintained by serial passage through Sprague-Dawley rats, as described previously(48). Third-stage larvae (L3) were isolated from the clean edges of parasite cultures and washed five times with PBS (Dulbecco’s PBS, Sigma) before infection. On day 0, mice were infected percutaneously with 500 L3 larvae on the right ear, as described in the results section. At various time points mice were euthanised and BAL and lungs were taken for further analysis. On day 1 p.i., BAL fluid absorbance was measured at 400nm using a VersaMax microplate reader (Molecular Devices) to assess lung haemorrhage(49). For lung-stage L4 counts on days 1 and 2 p.i., lungs were minced and incubated in PBS for 24h at 37°C; emergent larvae were counted using a dissecting microscope. For gut worm counts, small intestines of infected mice were collected in PBS, and were then cut longitudinally along the entire length, placed in gauze in a 50ml Falcon and incubated at 37°C for 3 to 4h, settled worms were then counted with the aid of a dissecting microscope.

### Histology and immunofluorescence staining

For histology, ears were collected in PBS and fixed in formalin for 24h before transferring to 70% ethanol. Tissues were processed, embedded in paraffin, and then sectioned (5 µm thickness) onto glass microscopy slides (Superfrost Plus Adhesion slides, J1830AMNZ, Thermo Fisher Scientific). For immunostaining, lung sections were deparaffinised, rehydrated and heat-mediated antigen retrieval performed using Tris-EDTA buffer (10 mM Tris, 1 mM EDTA), pH 9.0 (20 mins incubation at 95°C). Tissue sections were permeabilised in 1% Triton-X100 in PBS (10 mins at room temperature) and then non-specific protein-binding blocked (10% normal donkey serum, 1% BSA, 0.05% Tween-20 in PBS, 30 minutes at room temperature) before sections were incubated with primary antibody overnight at 4°C. All sections were washed in PBS with 0.05% Tween-20 followed by PBS before incubation with secondary antibody (donkey anti-rabbit NorthernLights™ NL637-conjugated antibody, 1:200, NL005; R&D Systems, or streptavidin NL557-conjugated, 1:800, NL999, R&D Systems) for 1 hour at room temperature. Sections were washed as before, incu-bated with 4,6-diamidino-2-phenylindole (DAPI, 1:40,000 in PBS, 2 mins at room temperature) and then mounted (Fluorescence mounting medium, S3023, Agilent Technologies) and cover-slipped. Staining was visualised on an Olympus IX83 deconvolution microscope.

### Cell preparation, flow cytometry, and analysis

BAL was collected in a 15mL Falcon tube by washing the airways of euthanised mice 4 times with 0.4 mL of PBS. Cells were centrifuged at 400 x g for 5 mins, the supernatant was either discarded or collected for further analysis; cells were then resuspended in PBS containing 1% FBS and 0.2 µM EDTA. Ears were dissected, and the two ear sheets were split with the help of a pair of forceps. Split ear sheets were then digested with Liberase (0.25 mg/mL; Roche) and DNase I (0.5 mg/mL; Sigma-Aldrich) in 0.5mL RPMI 1640 at 37°C, 200 rpm, for 1h 45 mins. The supernatant and the remainder of the tissue were added to a Medicon (BD Biosciences) then homogenised with the help of a Medimachine for 5 minutes. The resulting tissue homogenate was then passed through a 70-µm cell strainer and washed in 10mL of RPMI 1640 containing 10% Fetal Bovine Serum (FBS). Lungs were dissected and tissue was cut into pieces, then digested with Liberase (0.4 U/mL; Roche) and DNase I (80 U/ml) (Thermofisher) in RPMI 1640 at 37°C for 45 mins. Single cells were isolated by passing the tissue through a 70-µm cell strainer. Cells were washed in 10mL of RPMI 1640 containing 10% FBS. For lung samples, red blood cells were lysed using an ammonium-chloride-potassium (ACK) buffer (Gibco) for 3mins at room temperature and reaction was stopped by diluting samples in PBS. Total BAL, ear and lung live cells were counted with AO/PI dye on an automated cell counter (Auto2000, Nexcelom). Cells were then incubated with anti-CD16/CD32 5 µg/ml (Mouse Fc Block, BD) and LIVE/DEAD Blue (ThermoFisher) in PBS for 15 mins. Next, cells were incubated, without washing, with antibodies to surface antigens for further 15 minutes. After incubation cells were washed with PBS containing 1% FBS and 0.2 µM of EDTA and fixed with eBioscienceTM IC Fixation Buffer (Invitrogen) – or BD Cytofix™ Fixation Buffer (BD Biosciences) for cells expressing fluorescent proteins – for 20 minutes. Optionally, for intracellular cytokine staining, after tissue digestion and cell counting, cells were incubated in a 96 round-bottom wells plate in a 2-3x10^6^ cells/well concentration with a cell stimulation cocktail containing protein transport inhibitor (eBioscience) for 3h to 4h at 37°C. Cells were then stained for surface markers as described above, fixed for 30 mins at 4°C and permeabilised with the Foxp3/Transcription Factor Staining Buffer set (eBioscience) – or fixed with BD Cytofix™ Fixation Buffer (BD Biosciences) for cells expressing fluorescent proteins – and finally incubated overnight at 4°C with cytokine-specific antibodies in permeabilization buffer (eBioscience). Cells were analysed using either a FACSFortessa or a FACSymphony (both BD Biosciences) and analysed using FlowJo software (v10; Tree Star).

### Immuno-detection of cytokines in lung tissue samples

Ear tissue was homogenised in 2 mL sterile tubes containing 600 µL PBS plus a cocktail of protease inhibitors (cOmplete ULTRA tablets, Mini; Roche) and metal beads (BioSpec Products), with the help of a TissueLyzer II tissue homogenizer (Qiagen). The homogenate was centrifuged at 10000 x g to remove cellular debris and beads and stored at -80ºC until usage. Cytokine concentrations were assessed using LEGENDplex™ Mouse Th1/Th2 and Mouse Cytokine Panel 2 kits – both bead-based assays that use the principles of sandwich ELISA to quantify soluble analytes using a flow cytometer (Biolegend) – according to manufacturer’s instruction. Cytokine concentrations were normalised using total protein concentration in tissue homogenate, determined in a NanoDrop spectrophotometer (ThermoFisher).

### Nanostring RNA profiling

Lung and ear tissue were homogenised in 2 mL sterile tubes containing 300 µL of lysis buffer (Qiagen), with the help of a TissueLyzer II tissue homogenizer (Qiagen). mRNA was then prepared from tissue homogenates using the RNeasy^®^ Mini Kit (Qiagen). Extracted RNA was then diluted to 20ng/µL in RNase free H2O, measured using Qubit™ RNA HS Assay Kit (Thermofisher) and run on a Nanostring nCounter^®^ FLEX system using the Mouse Host Response (V1) and Mouse Myeloid Innate Immunity (V2) panels as specified in legends as per manufacturer’s instructions. Raw data were loaded into nSolver version 4.0 using default settings. Non-normalised counts were then exported, and subsequent analyses were performed in R (v3.6) using RStudio (v1.2.1335 Build 1379). Positive controls were analysed to ensure there was clear resolution at variable expression levels and negative controls were used to set a minimum detection threshold which was applied to all samples. Data were then normalised with edgeR (v4.0) using the TMM method and differential expression between *N. brasiliensis*-infected and naïve mice, or p.c.- and i.v.-infected mice, was calculated via linear modelling with empirical Bayes smoothing using the limma R package (v2.43). Genes with an absolute fold change of greater than one and a significance value of under 0.05 were defined as “differentially expressed” and taken forward for further analysis. PCA and Volcano plots were then generated from normalised counts and fold changes of DEGs using the ggplot2 R package (v3.5.1). The networks and functional analyses of DEGs were generated either via the clusterProfiler (v4.10.1) and enrichplot (v1.22.0) packages, or using the Ingenuity Pathway Analysis software (IPA; QIAGEN Inc., https://www.qiagenbio-informatics.com/products/ingenuity-pathway-analysis); for the latter, data were then imported into R for visualisation.

### Cell sorting

For cell sorting of lung innate-like lymphocytes, lung tissue was digested as described above and, additionally, single cells were isolated by passing the tissue through a 40-µm cell strainer, followed by a 70% / 30% Percoll (Sigma-Aldrich) gradient and 30-min centrifugation at 2400 rpm. Leukocytes were recovered from the interface, resuspended, and immunostained as described above. Up to 10,000 γδ T cells and Lin^-^Thy1.2^+^ cells (defined as CD45^+^CD64^-^CD11b^-^CD11c^-^MHC-II^-^B220^-^Thy1.2^+^CD4^-^γδTCR^+/-^), 2,500 NK1.1^+^γδTCR^-^ innate lymphocytes (defined as CD45^+^CD64^-^CD11b^-^CD11c^-^MHC-II^-^B220^-^Thy1.2^+^CD4^-^γδTCR^-^), and 2,500 NK cells (defined as CD45^+^Thy1.2^-^NK1.1^+^) per mouse were then sorted on a FACSAria (BD Biosciences) into sterile RNase-free 2 mL tubes containing RPMI 1640 with 10% FBS. The cell populations sorted from 4 mice per group were then pooled post-sorting in one Eppendorf containing PBS with 10

### Single cell isolation and library construction

Gene expression libraries were prepared from single cells using the Chromium Controller and Single Cell 3 Reagent Kits v3.1 (10x Genomics, Inc. Pleasanton, USA) according to the manufacturer’s protocol (CG000315 Rev C). Briefly, nanoliter-scale Gel Beads-in-emulsion (GEMs) were generated by combining barcoded Gel Beads, a master mix containing cells, and partitioning oil onto a Chromium chip. Cells were delivered at a limiting dilution, such that the majority (90-99%) of generated GEMs contain no cell, while the remainder largely contain a single cell. The Gel Beads were then dissolved, primers released, and any co-partitioned cells lysed. Primers containing an Illumina TruSeq Read 1 sequencing primer, a 16-nucleotide 10x Barcode, a 10-nucleotide unique molecular identifier (UMI) and a 30-nucleotide poly(dT) sequence were then mixed with the cell lysate and a master mix containing reverse transcription (RT) reagents. Incubation of the GEMs then yielded barcoded cDNA from poly-adenylated mRNA. Following incubation, GEMs were broken and pooled fractions recovered. First-strand cDNA was then purified from the post GEM-RT reaction mixture using silane magnetic beads and amplified via PCR to generate sufficient mass for library construction. Enzymatic fragmentation and size selection were then used to optimize the cDNA amplicon size. Illumina P5 & P7 sequences, a sample index, and TruSeq Read 2 sequence were added via end repair, A-tailing, adaptor ligation, and PCR to yield final Illumina-compatible sequencing libraries.

### Single Cell RNA Sequencing

The resulting sequencing libraries comprised standard Illumina paired-end constructs flanked with P5 and P7 sequences. The 16 bp 10x Barcode and 10 bp UMI were encoded in Read 1, while Read 2 was used to sequence the cDNA fragment. Sample index sequences were incorporated as the i7 index read. Paired-end sequencing (28:90) was performed on the Illumina NovaSeq6000 platform.

### Data pre-processing

Raw data files generated from the sequencing instrument were processed using 10x Genomics Cell Ranger pipeline (v7.1.0). Base call (BCL) files were demultiplexed and converted to FASTQ files using “cellranger mkfastq”. The FASTQ files were then mapped to the pre-built Mouse reference package from 10x Genomics (mm10-2020-A) using “cellranger count” to produce the feature-barcode matrices.

### Single cell data analysis

The single-cell data were processed in R environment (v4.3) following the workflow documented in Orchestrating Single-Cell Analysis with Bioconductor(50). Briefly, for each sample, the HDF5 file generated by Cell Ranger was imported into R to create a SingleCellExperiment object. A combination of median absolute deviation (MAD), as implemented by the “isOutlier” function in the scuttle R package (v1.12.0) and exact thresholds was used to identify and subsequently remove low quality cells before data integration. The log-normalised expression values of the combined data were computed using the “multiBatchNorm” function from the batchelor R package (v1.18.1). The per-gene variance of the log-expression profile was modelled using the “modelGeneVarByPoisson” function from the scran R package (v1.30.2) and top 5,000 highly variable genes were selected. The mutual nearest neighbors (MNN) approach implemented by the “fastMNN” function from the batchelor R package was used to perform batch correction. The first 50 dimensions of the MNN low-dimensional corrected coordinates for all cells were used as input to produce the t-stochastic neighbour embedding (t-SNE) projection and uniform manifold approximation and projection (UMAP) using the “runTSNE” and “runUMAP” functions from the scater R package (v1.30.1) respectively. The Leiden algorithm from the igraph R package (v2.0.2) was used to perform a first-pass clustering, followed by clustering using the Louvain algorithm to produce 39 putative cell clusters. Through manual curation, the cells were grouped into 20 clusters, including one consisting of doublet cells. Differential expression analysis of the 3 treatment groups (control, *N. brasiliensis* infection via skin contact and infected intravenously) across clusters was performed using DESeq2 R package (v 1.43.4), with recommended parameters. Genes with a false discovery rate (FDR) below 5% were considered differentially expressed.

To calculate a “Type-1 module score”, the combined per-cell mean expression of candidate type-1 effector molecules extracted from the literature(27–29) was calculated using the “colMeans” function in the gmatrix R package.

### Bulk RNA sequencing

For cell sorting of dermal γδ T cells and DETCs, skin tissue was digested as described above and, additionally, single cells were isolated by passing the tissue through a 40-µm cell strainer, followed by a 70% / 30% Percoll (Sigma-Aldrich) gradient and 30-min centrifugation at 2400 rpm. Leukocytes were recovered from the interface, resuspended, and immunostained as described above. Dermal γδ T cells were defined as CD45^+^CD64^-^CD11b^-^CD11c^-^MHC-II^-^FcεR1^-^Thy1.2^+^CD3^+^γδTCR^+^Vγ5^-^, DETCs were defined as CD45^+^CD64^-^CD11b^-^CD11c^-^MHC-II^-^FcεR1^-^Thy1.2^+^CD3^+^γδTCR^+^Vγ5^+^ and then electronically sorted on a FACSAria (BD Biosciences) into sterile RNase-free Eppendorfs containing 200µL of lysis buffer (Qiagen), for posterior mRNA isolation; each mouse was treated as one individual sample. RNA was then isolated from sorted cells using the Single Cell RNA Purification Kit (Norgen). Samples were sent to Novogene Company Limited (Cambridge, UK) where QC checks and sequencing were performed. mRNA pre-amplification was employed using the SMARTer kit (Takara Bio) prior to library preparation by poly(A) enrichment and paired-end sequenced on an Illumina NovaSeq 6000 for 300 cycles (2 x 150 bp). FASTQ files were received, which were then aligned to GENCODE mouse genome (version M33, Jan 2020), and quantified using Salmon software (v1.10.3) using –validatemappings and –gcBias flags (COMBINE-lab). Annotated and quantified data was then imported into RStudio (v2022.07.2+576) using the Tximeta R package (v4.2.3). Genes with an average raw count of less than ten across all groups were removed. Principal component analysis (PCA) was used to summarise total variation within the dataset. Differential expression analysis was performed using the DESeq2 R package (v1.46.0) and volcano plots and dot plots were then generated using ggplot2 R package (v3.5.1). The networks and functional analyses of DEGs were generated either via the clusterProfiler (v4.10.1) and enrichplot (v1.22.0) packages or via import into PANTHER (v19.0) via the web GUI (https://geneontology.org).

### In vivo *substance administration*

4-hydroxytamoxifen (4OH-T; Sigma-Aldrich) was diluted in 99% ethanol to a final concentration of 12 mg/mL. Experimental mice had their ears shaved 1 day prior to the start of treatment; 15 µL was then applied on the ears of previously anaesthetised mice (10 µL on the external side and 5 µL on the inner side) for 5 consecutive days. Animals were then rested for 3 days before being used for N. brasiliensis infection or ex vivo analysis.

For FTY720 experiments, FTY720 (Sigma-Aldrich) was administered in the drinking water (final concentration of 2.5 µg/mL) three days before infection. Water pouches were replaced after three days with freshly prepared FTY720 added each time.

For IL-1R blockade experiments mice were treated s.c. with 100 mg/Kg of Anakinra (Kineret; Swedish Orphan Biovitrum AB) 30 minutes p.i.; a second dose was given at 3.5 hours p.i. in all the experiments where mice were analysed at 6h p.i. and in one of the experiments where mice were analysed at 24h p.i. Control mice received the same volume of a placebo formulation (Swedish Orphan Biovitrum AB) via the s.c. route.

### Skin photo-labelling

Kaede mice had their ear skin photo-labelled using a 405-nm wavelength focused LED light (Dymax BlueWave QX4 fitted with 8 mm focusing lens, DYM41572; Intertronics) as described previously(51).

### Statistics

Prism 9.5.1 and 10.4.0 (versions 9 and 10, respectively; GraphPad Software Inc) were used for statistical analysis. Data were tested for normality using the Shapiro-Wilk test and then differences between experimental groups were assessed accordingly. For gene expression data, normalised expression values were log2 transformed to achieve normal distribution. Comparisons with a *p* value<0.05 (unless stated otherwise) were considered to be statistically significant. Data are represented as mean±SD.

## ACKNOWLEDGEMENTS

We thank the staff at the Biological Services Facility, Flow Cytometry Facility and Genomics Facility – in particular David Chapman and Claire Morrisroe – at The University of Manchester. We are also grateful to Novogene for the services provided. We thank Miguel Muñoz-Ruiz, Kara Filbey, Jesuthas Ajendra, Alistair Chenery, Conor Finlay, Tara Sutherland, Dominik Rückerl, Suzie Hodge, Roser Tachó-Piñot, Kelly Wemyss, Stuart Allan, David Bending, Juan Quintana, Amy Saunders, Lizzie Mann and Abdullah Shazad for helpful discussions, sharing of resources and technical advice. Finally, we thank Prof. Bruno Silva-Santos for carefully reading the manuscript. This work was funded by the Wellcome Trust (106898/A/15/Z) and MRC (MR/V011235/1) to J.E.A., European Molecular Biology Organization (LTF 191-2019) and European Commission Marie Sklodowska-Curie Individual Fellowship (ref. 101025781) to P.H.P.; Sir Henry Dale Fellowship (jointly funded by the Wellcome Trust and the Royal Society; 105644/Z/14/Z) and BBSRC (BB/T014482/1) to M.R.H.

## AUTHOR CONTRIBUTIONS

Conceptualization: P.H.P. and J.E.A.; Methodology and Investigation: P.H.P., J.E.P., B.H.K.C., G.E.B., F.D., L.Z., R.J.D., S.T.P., N.F., M.R.H.; Data Analysis: P.H.P., J.E.P. and I-H.L.; Resources: D.R.W. and J.E.K.; Writing – original draft: P.H.P. and J.E.A.; Writing – review and editing: P.H.P., J.E.P., J.E.K., M.R.H., and J.E.A.

## COMPETING FINANCIAL INTERESTS

The authors declare no conflicts of interest.

## Supplementary Information

**Extended Data Fig. 1.**
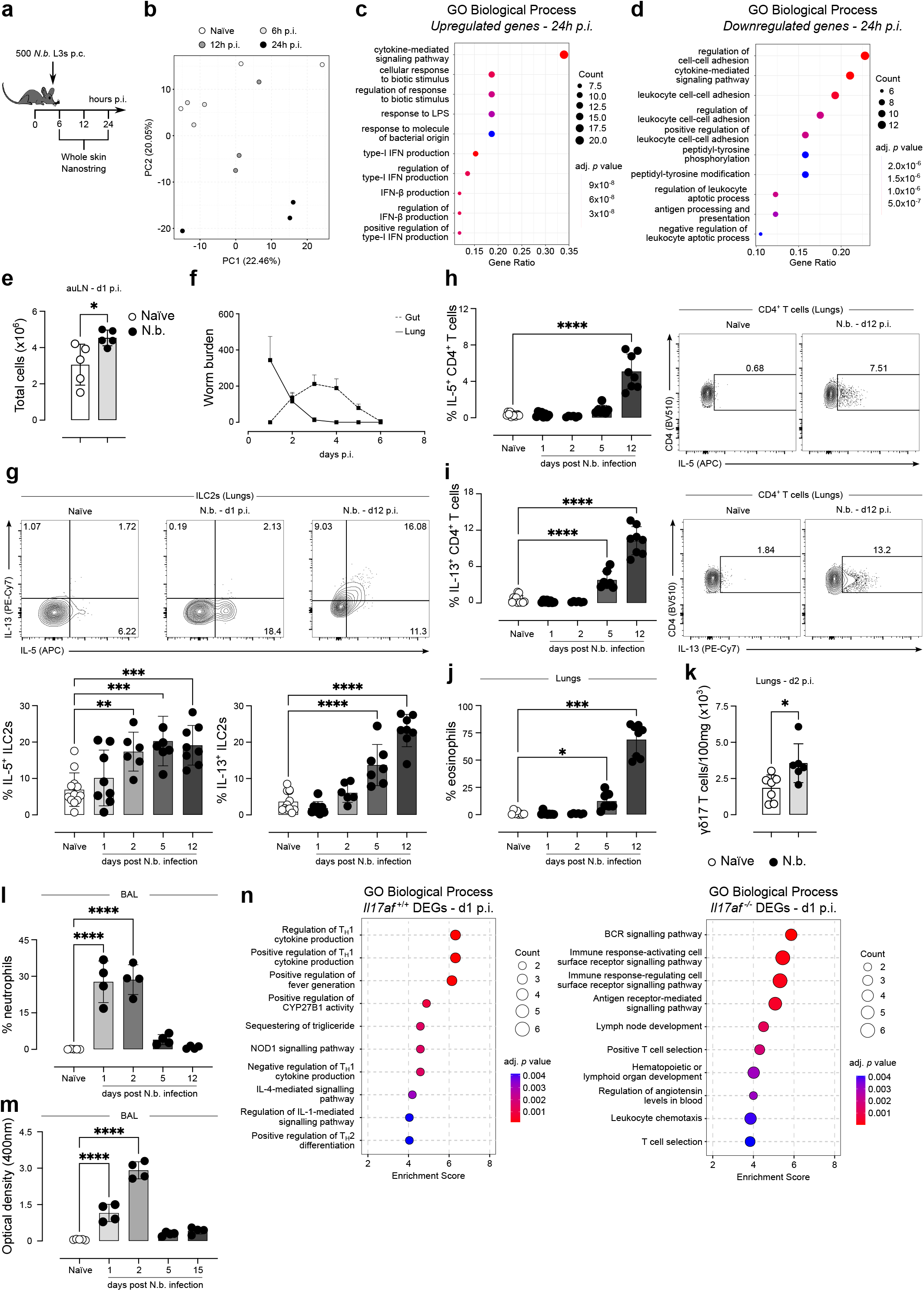
*N. brasiliensis* p.c. infection recapitulates the hallmarks of the s.c. route of infection. **a**, Mice were infected with 500 L3s as described in Fig. 1a and ears were collected at 6h, 12h and 24h p.i. for subsequent NanoString analysis. **b**, Skin RNA from naïve (white circles), and *N. brasiliensis*-infected (light grey to black circles) C57BL/6J mice was analysed with the Mouse Host Response V1 NanoString panel and total sample variation was summarised via principal component analysis (PCA) across all genes. (*n* = 3 mice per group). **c, d**, Overrepresented GO: biological processes in upregulated **(c)** and downregulated **(d)** DEGs from *N. brasiliensis*-infected at 24h p.i. **e**, Summary graphs showing the cell numbers in the skin-draining lymph nodes in naïve (circles) and *N. brasiliensis* infected (triangles) C57BL/6J mice (*n* = 5 mice per group; representative of at least 3 independent experiments). **f**, worm burden measured in the lungs (full line) and small intestines (dashed line) of mice infected s.c. with 500 *N. brasiliensis* L3s throughout time. **g**, Flow cytometry analysis of intracellular IL-5 and IL-13 in ILC2s isolated from the lungs of naïve and *N. brasiliensis*-infected C57BL/6J mice and stimulated ex vivo for 4h. Summary graph showing the percentages of IL-5^+^ and IL-13^+^ cells within the ILC2 population (*n* = 6-12 mice per group; pool of 2 independent experiments). **h, i**, Flow cytometry analysis and summary graphs of intracellular IL-5 **(h)** and IL-13 **(i)** in CD4^+^ T cells isolated from the lungs of naïve and *N. brasiliensis*-infected C57BL/6J mice and stimulated ex vivo for 4h (*n* = 6-12 mice per group; pool of 2 independent experiments). **j**, Frequencies of eosinophils within total leukocytes in the BAL throughout the course of *N. brasiliensis* p.c. infection (*n* = 6-9 mice per group; pool of 2 independent experiments). **k**, Numbers of γδ17 T cells per 100 mg of tissue isolated from the lungs of naïve (white circles) and *N. brasiliensis*-infected C57BL/6J mice (black circles) at d2 p.i. (*n*= 6-7 mice; pool of 2 independent experiments for). **l, m**, Frequencies of neutrophils within total leukocytes **(l)**, and optical density (measured at 400nm) **(m)** in the BAL throughout the course of *N. brasiliensis* s.c. infection (500 *N. brasiliensis* L3s; *n* = 4-5 mice per group). **n**, Lung RNA from naïve and *N. brasiliensis*-infected *Il17a/f* ^+/+^ and *Il17a/f* ^-/-^ mice was analysed with the Mouse Host Response V1 NanoString panel. Overrepresented GO: biological processes in *Il17a/f* ^+/+^ (left) and and *Il17a/f* ^-/-^ (right) DEGs (naïve vs. *N. brasiliensis*-infected) at 24h p.i. (*n* = 3 mice per group). e-m, Error bars represent mean±SD. Normality of the samples was assessed with Shapiro-Wilk normality test; statistical analysis was then performed accordingly. * *p* < 0.05; ** *p* < 0.01; **** *p* < 0.0001

**Extended Data Fig. 2.**
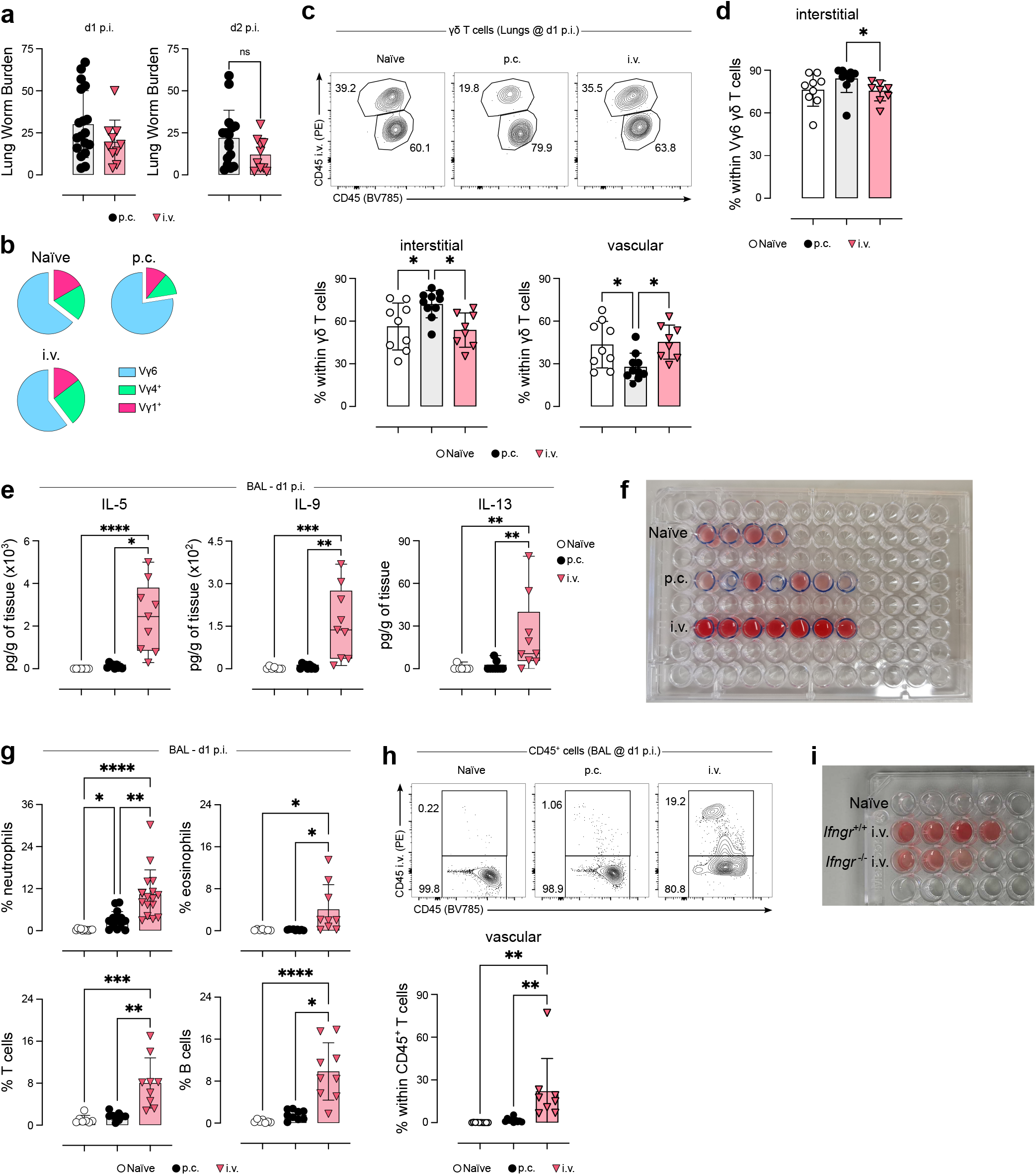
Bypassing the skin phase of infection precludes the induction of lung type-17 responses and results in increased IFN-γ-dependent lung bleeding during *N. brasiliensis* larvae migration. **a**, Worm burden measured in the lungs of mice infected p.c. with 500 *N. brasiliensis* L3s (black circles) and i.v. with with 200 *N. brasiliensis* L3s (pink triangles), at days 1 and 2 p.i.. **b**, Pie charts depicting the Vγ chain usage among lung γδ T cell in naïve, p.c., and i.v. *N. brasiliensis*-infected C57BL/6J mice (*n* = 8-10 mice per group; pool of 2 independent experiments). c, d, C57Bl6/J mice were infected percutaneously with 500 *N. brasiliensis* L3s; for the i.v. route, mice were injected with 200 L3s in sterile PBS; the naïve group was left uninfected; before being culled at day 1 p.i. all mice were injected i.v. with a fluorochrome-labelled CD45 antibody. **c**, Summary graphs showing the frequencies of interstitial (left) and vascular (right) γδ T cells; representative FACS plots depicting the staining with CD45 i.v. and CD45 in the antibody mix (*n* = 8-10 mice per group; representative of 2 independent experiments). **d**, Summary graphs showing the frequencies of Vγ6 cells (defined as Vγ1^-^Vγ4^-^) cells within the γδ T cell population in naïve (white circles), p.c.-infected (black circles), and i.v.-infected (pink triangles) mice (*n* = 8-10 mice per group; representative of 2 independent experiments). **e**, Cytokine levels in total lung tissue from naïve (white circles), p.c. (black circles) and i.v. (pink triangles) *N. brasiliensis*-infected C57BL/6J mice normalised by 1 gram of tissue, measured by LEGENDPlexTM (*n* = 7-9 mice per group; pool of 2 independent experiments). **f**, Representative picture of the measurements shown in Fig. 2i (*n* = 4-7 mice per group). **g**, Frequencies of neutrophils, eosinophils, T cells and B cells within total leukocytes in the lungs of naïve (white circles), p.c. (black circles) and i.v. (pink triangles) *N. brasiliensis*-infected C57BL/6J mice at day 1 p.i. (*n* = 7-9 mice per group; pool of 2 independent experiments). **h**, C57Bl6/J mice were infected percutaneously with 500 *N. brasiliensis* L3s; for the i.v. route, mice were injected with 200 L3s in sterile PBS; the naïve group was left uninfected; before being culled at day 1 p.i. mice from all groups were injected i.v. with a fluorochrome-labelled CD45 antibody. Summary graph showing the frequencies of vascular leukocytes in the BAL (bottom). Representative FACS plots depicting the staining with CD45 i.v. and CD45 in the antibody mix in BAL cells (top) (*n* = 8-10 mice per group; representative of 2 independent experiments). **i**, Representative picture of the measurements shown in Fig. 2j (*n* = 3-4 mice per group). **a-h**, Error bars represent mean±SD. **e**, Box-and-whisker plots display first and third quartiles, and the median; whiskers are from each quartile to the minimum or maximum. **a-h**, Normality of the samples was assessed with Shapiro-Wilk normality test and statistical analysis was then performed accordingly. * *p* < 0.05; ** *p* < 0.01; *** *p* < 0.001; **** *p* < 0.0001

**Extended Data Fig. 3.**
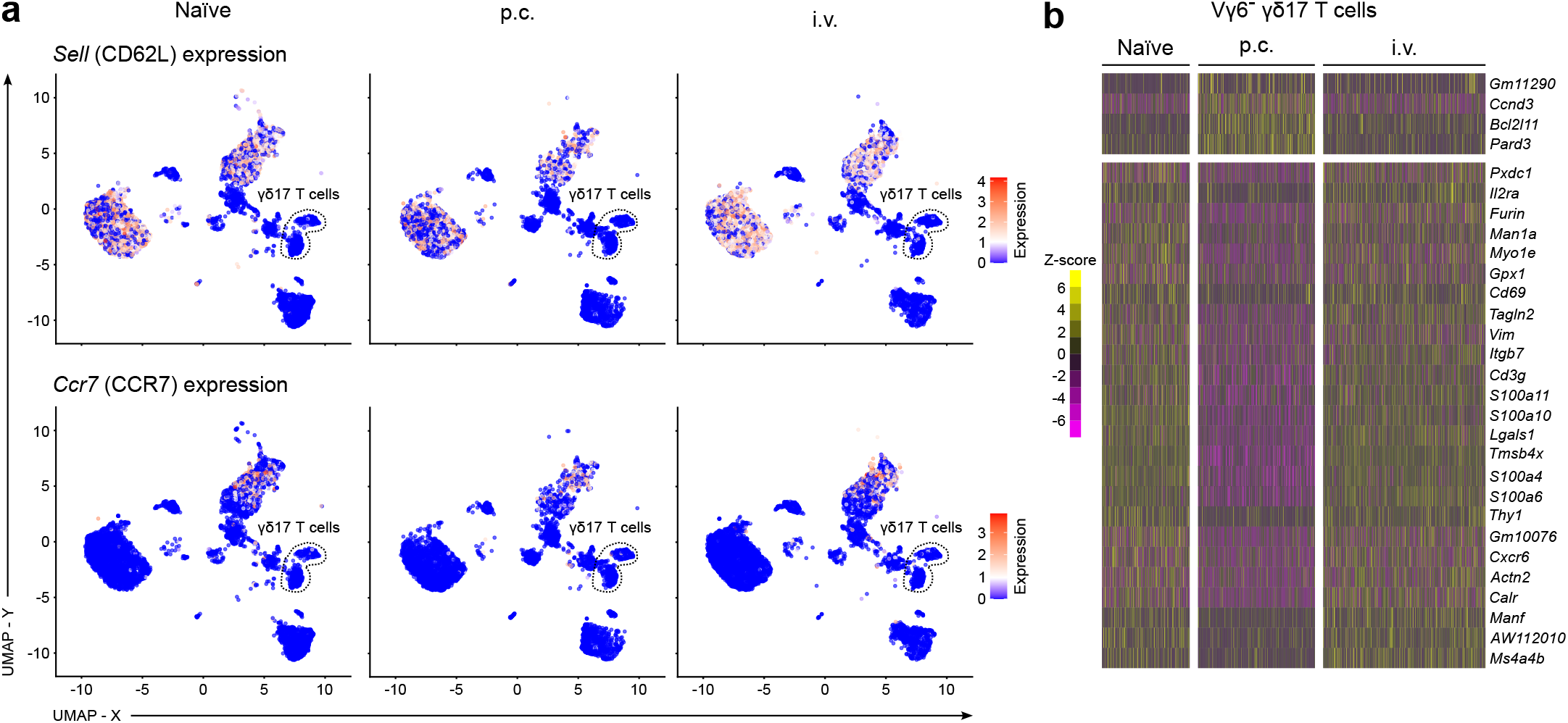
P.c., but not i.v., *N. brasiliensis* infection induces the activation of Vγ6^+^ γδ17 T cells and suppression of IFN-γ-producing innate lymphocytes. **a**, *Sell* and *Ccr7* expression projected onto two dimensional UMAP plot and separated by infection status. γδ T cell populations are shown in the dotted area. **b**, Heatmap showing the DEGs in Vγ6^-^ γδ17 T cells from naïve, p.c.- and i.v.-infected animals. Scale represents scaled and centred z-scores. scRNAseq data in **(a)** and **(b)** consists of sorted cells pooled from 4 mice per group.

**Extended Data Fig. 4.**
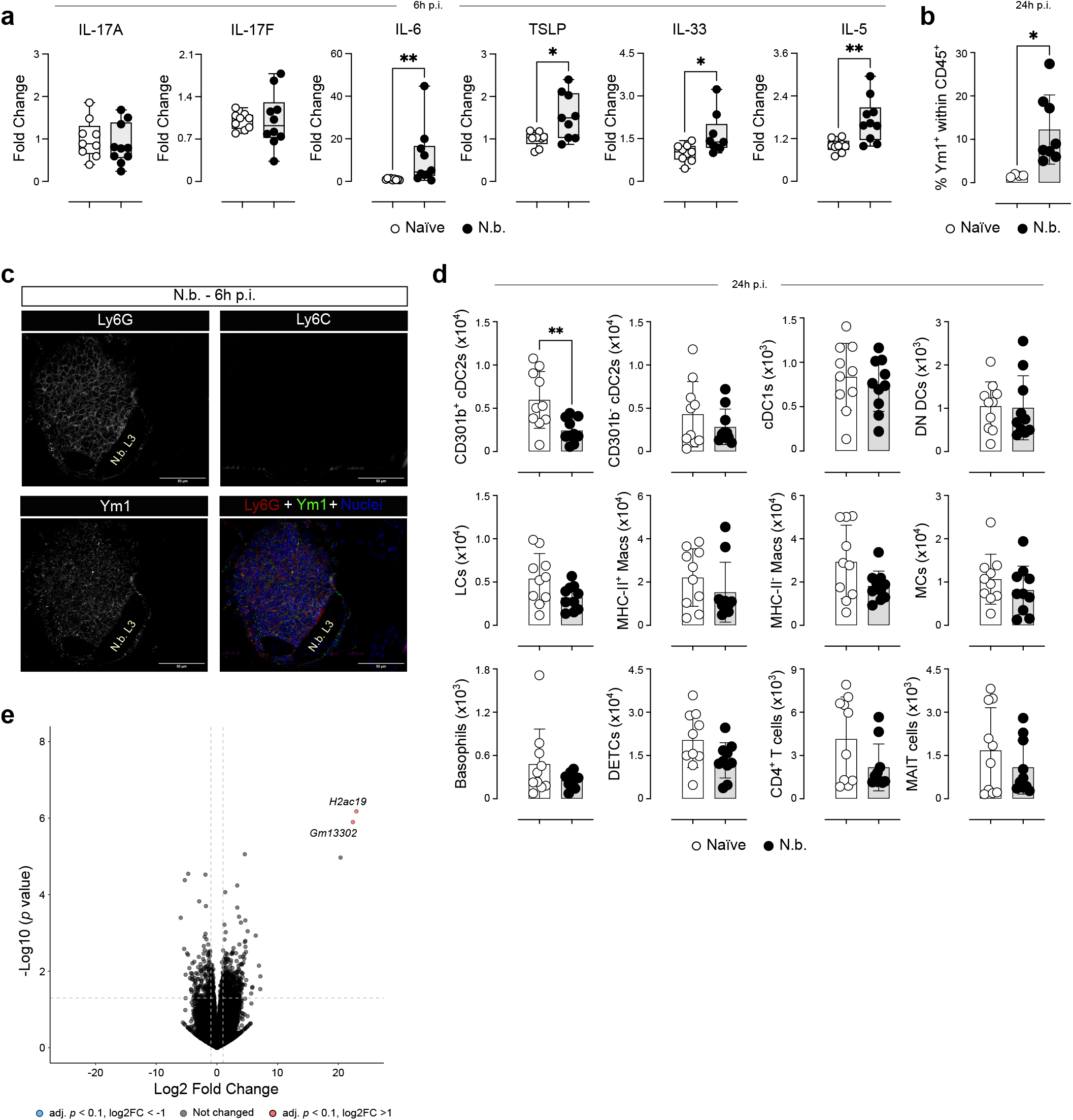
*N. brasiliensis* larvae induce a migratory transcriptional program in dermal γδ T cells. **a**, Cytokine levels in total skin tissue from naïve (white circles), and *N. brasiliensis*-infected (black circles) C57BL/6J mice measured by LEGENDPlex™ and displayed as fold change in relation to the average of the naïve group (n = 9-10 mice per group; pool of 2 independent experiments). **b**, Summary graph from the flow cytometry analysis of intracellular Ym1 in total leukocytes isolated from the skin of naïve (white circles) and *N. brasiliensis*-infected (black circles) mice (n = 4-8 mice per group; pool of 2 independent experiments). **c**, Immunofluorescence of *N. brasiliensis* larvae and recruited immune cells in the skin of C57BL/6J mice at 6h p.i. **d**, Summary graphs showing the numbers of skin immune cell populations from naïve (white circles) and *N. brasiliensis*-infected (black circles) mice at 24h p.i. (n = 9-10 mice per group; pool of 2 independent experiments). **e**, DEGs from DETCs were visualised using a volcano plot, genes significantly upregulated (red), and downregulated (blue) during *N. brasiliensis* infection were defined as having an adjusted *p* value of <0.1 and absolute log2 fold change of >1. a, Box-and-whisker plots display first and third quartiles, and the median; whiskers are from each quartile to the minimum or maximum. **b**,**d**, Error bars represent mean±SD. **a, b, d**. Normality of the samples was assessed with Shapiro-Wilk normality test and statistical analysis was then performed accordingly. * *p* < 0.05; ** *p* < 0.01.

**Extended Data Fig. 5.**
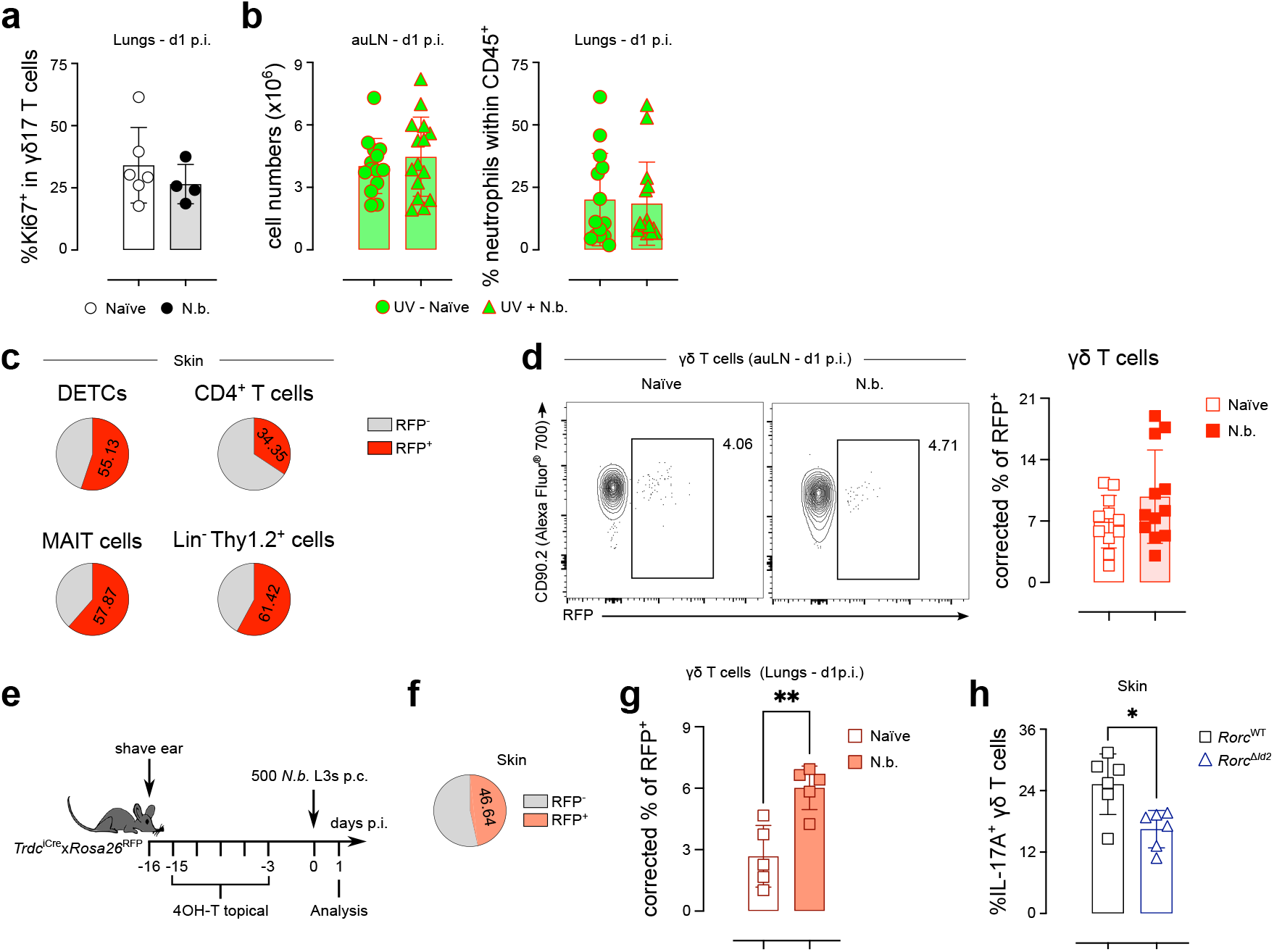
Migratory dermal γδ17 T cells are responsible for the expansion in the lung γδ17 T cell population during *N. brasiliensis* infection. **a**, Summary graph from the flow cytometry analysis of intracellular Ki67 in γδ17 T cells isolated from the lungs of naïve (white circles) and *N. brasiliensis*-infected (black circles) mice (n = 4-6 mice per group). **b**, Summary graphs showing the cell numbers in the skin-draining lymph nodes (left) and frequencies of neutrophils in the lungs (right) of naïve (circles) and N. brasiliensis infected (triangles) photo-labelled Kaede mice (n = 14-15 mice per group; pool of 3 independent experiments). **c**, Summary pie charts showing the percentage of RFP^+^ cells (red slice) within DETCs, CD4^+^ T cells, MAIT cells and Lin-Thy1.2^+^ cells in naïve *Id2*^CreERT2^x*Rosa26*^RFP^ mice (n = 11-13 mice). **d**, Flow cytometry analysis of RFP expression and summary graph showing the corrected percentage of RFP^+^ cells within total γδ T cells isolated from the ear-draining lymph nodes of naïve (white squares) and *N. brasiliensis*-infected (red squares) *Id2*^CreERT2^x*Rosa26*^RFP^ mice (n = 11-12 mice per group; pool of 3 independent experiments). **e**, *Trdc*^CreERT2^x*Rosa26*^RFP^ mice had their right ear shaved, and starting on the following day were treated for five times with 4OH-T over the course of two weeks. Mice were then rested for 3 days before being infected with 500 *N. brasiliensis* L3s p.c. **f**, Summary pie chart showing the percentage of RFP^+^ cells (orange slice) within dermal γδ T cells in the skin of naïve *Trdc*^CreERT2^x*Rosa26*^RFP^ mice (n = 10). **g**, Summary graph showing the percentages of RFP^+^ cells within γδ T cells isolated from the lungs of naïve (white squares) and *N. brasiliensis*-infected (orange squares) *Trdc*^CreERT2^x*Rosa26*^RFP^ mice (n = 5 mice per group). **h**, Summary graph showing the percentages of IL-17A^+^ cells within the dermal γδ T cell population isolated from the skin of naïve (white squares) and *N. brasiliensis*-infected (white triangles) *Id2*^CreERT2^x*Rorc*^fl/fl^ mice (n = 5-6 mice per group). **d, g**, Individual percentages of RFP^+^ cells were normalised to the respective Cre penetrance calculated from skin percentages of RFP^+^ cells, assuming 100% efficiency. **a-h**, Error bars represent mean±SD. Normality of the samples was assessed with Shapiro-Wilk normality test and statistical analysis was then performed accordingly. * *p* < 0.05.

**Extended Data Fig. 6.**
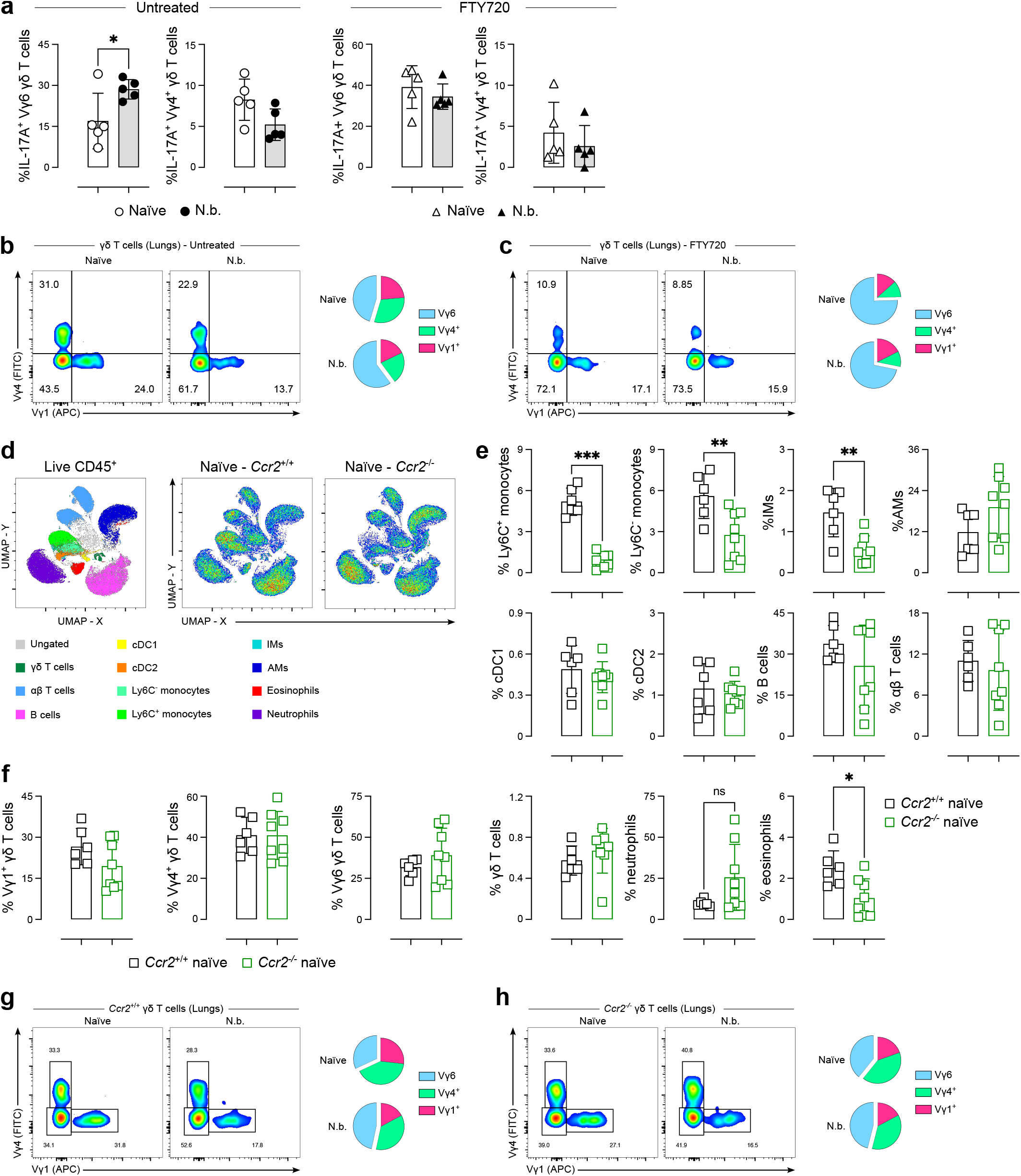
Dermal γδ17 T cells migrate to the lungs during *N. brasiliensis* infection in a S1PR- and CCR2-dependent fashion. **a**, Summary graphs showing the percentages of IL-17A^+^ cells within lung Vγ6 (defined as Vγ1^-^Vγ4^-^) and Vγ4^+^ γδ T cells from untreated and FTY720-treated naïve (white symbols) and *N. brasiliensis*-infected (black symbols) mice after 4h of ex vivo stimulation (*n* = 5 mice per group; representative of 3 independent experiments). **b, c**, Summary pie charts showing the Vγ usage in total lung γδ T cells from untreated naïve and *N. brasiliensis*-infected mice **(b)** and FTY720-treated naïve and *N. brasiliensis*-infected mice **(c)** (*n* = 5 mice per group; representative of 3 independent experiments). **d-f**, Flow cytometry analysis showing the immune cell populations in the lungs of naïve *Ccr2*^+/+^ and *Ccr2*^-/-^ mice. Representative UMAP plot (pool of 3 mice per group) summarising the flow cytometry analysis of lung immune cell populations (left) and density UMAP plots summarising the data for *Ccr2*^+/+^ (middle) and *Ccr2*^-/-^ (right) mice **(d)**. Frequency of immune cell populations within live CD45^+^ leukocytes (e) (*n* = 6-8 mice per group; pool of 3 independent experiments). Frequency of γδ T cell subsets within total γδ T cells **(f)** (n = 6-8 mice per group; pool of 3 independent experiments). **g, h**, Flow cytometry analysis and summary pie charts showing the Vγ usage in total lung γδ T cells from *Ccr2*^+/+^ (g) and *Ccr2*^-/-^ (**h)** mice (*n* = 6-10 mice per group; pool of 3 independent experiments). **a-f**, Error bars represent mean±SD. Normality of the samples was assessed with Shapiro-Wilk normality test and statistical analysis was then performed accordingly. * *p* < 0.05; ** *p* < 0.01; *** *p* < 0.001.

**Extended Data Fig. 7.**
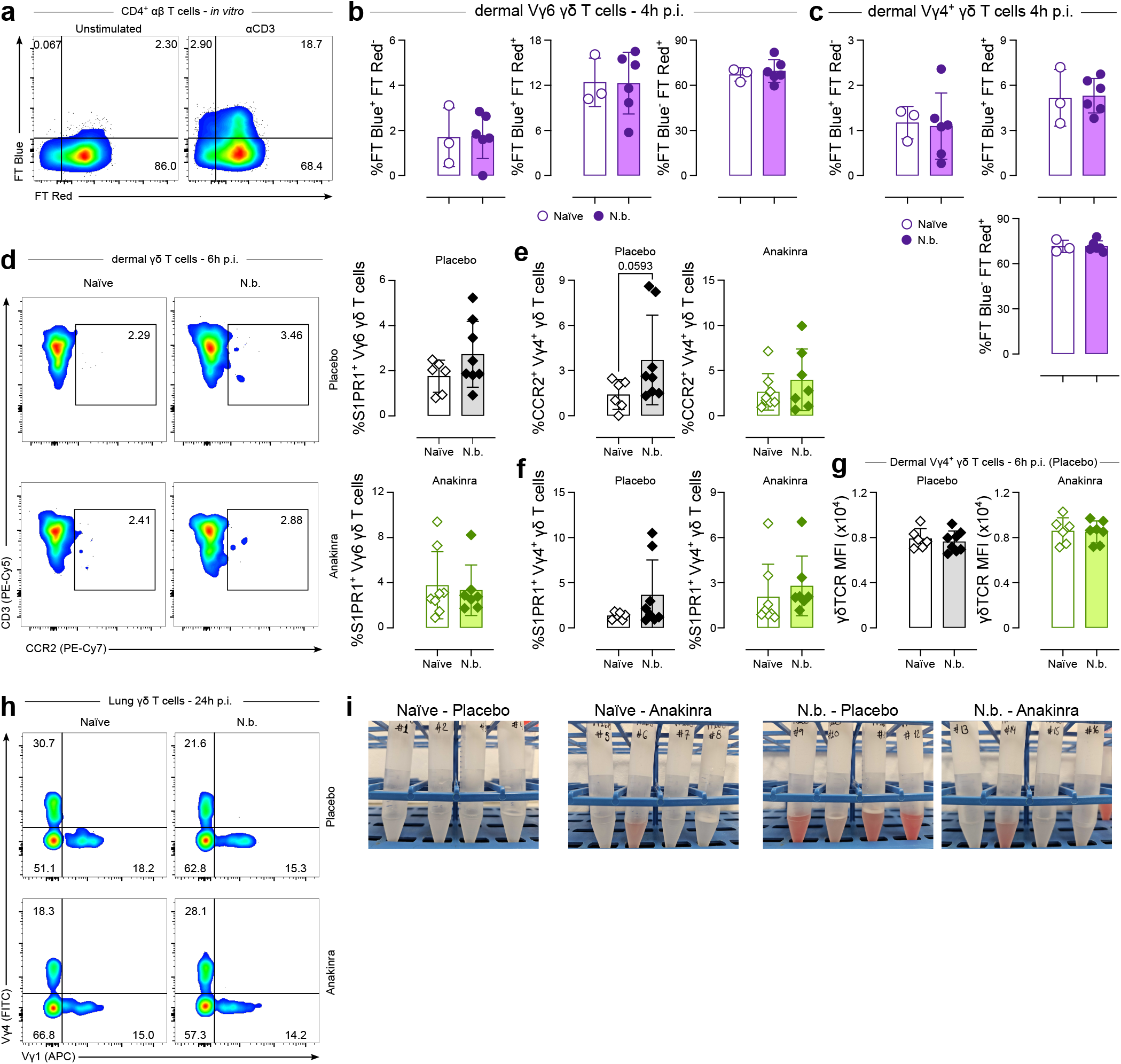
Dermal γδ T cells are activated via the IL-1R signalling pathway during *N. brasiliensis* infection. **a**, To validate the function of Nur77-Tempo mice, lymph node cells were stimulated in vitro with 2.5 µg/mL of anti-CD3 for 4h and FT Blue expression was analysed in CD4^+^ αβ T cells. were infected or not with 500 *N. brasiliensis* L3s p.c. and analysed at 4h p.i. **b, c**, Summary graphs showing FT protein expression among dermal Vγ6 (b) and Vγ4^+^ **(c)** γδ T cells from naïve (white circles) and *N. brasiliensis* infected (purple circles) mice (*n* = 3-6 mice per group; representative of 2 independent experiments). **d**, Flow cytometry analysis and summary graphs showing S1PR1 protein expression among dermal Vγ6 γδ T cells (defined as Vγ1^-^Vγ4^-^) from naïve (white diamonds) and *N. brasiliensis* infected (filled diamonds) mice treated with placebo (top) or Anakinra (bottom) (*n* = 6-8 mice per group; pool of 2 independent experiments). **e, f**, Summary graphs showing the percentage of CCR2-expressing **(e)**, and S1PR1-expresing **(f)** Vγ4^+^ γδ T cells from naïve (white diamonds) and *N. brasiliensis* infected (filled diamonds) mice treated with placebo (left) or Anakinra (right) (*n* = 6-8 mice per group; pool of 2 independent experiments). **g**, Summary graphs showing the median fluorescence intensity of surface γδTCR among dermal Vγ4^+^ γδ T cells from naïve (white diamonds) and *N. brasiliensis* infected (filled diamonds) mice treated with placebo (top) or Anakinra (bottom) (*n* = 6-8 mice per group; pool of 2 independent experiments). **h**, Flow cytometry analysis of the Vγ chain usage among lung γδ T cells from naïve and *N. brasiliensis* infected mice treated with placebo (top) or Anakinra (bottom) (*n* = 11-12 mice per group; pool of 3 independent experiments). **i**, Representative pictures of the data presented in Fig. 7i (*n* = 4 mice per group; representative of 3 independent experiments). Error bars represent mean±SD. Normality of the samples was assessed with Shapiro-Wilk normality test; statistical analysis was then performed accordingly **(b-g)**

**Extended Data Table 1.**

| **Type-1 effector genes** |
| --- |
| Ifng |
| Gzmb |
| Gzma |
| Cd7 |
| Tbx21 |
| Ccl5 |
| Klrc2 |
| Ly6c2 |
| Nkg7 |
| Klrd1 |
| Ctsw |
| Cd2 |
| Cd27 |
| Klre1 |
| Xcl1 |
| Lck |
| Cd28 |
| Tyrobp |
| Sell |
| Fasl |
| Ncr1 |
| Il2rb |
| Fcer1g |
| Itgb2 |
| Prf1 |
| Slamf7 |
| Runx3 |
| Lgals1 |

**Extended Data Table 2.**
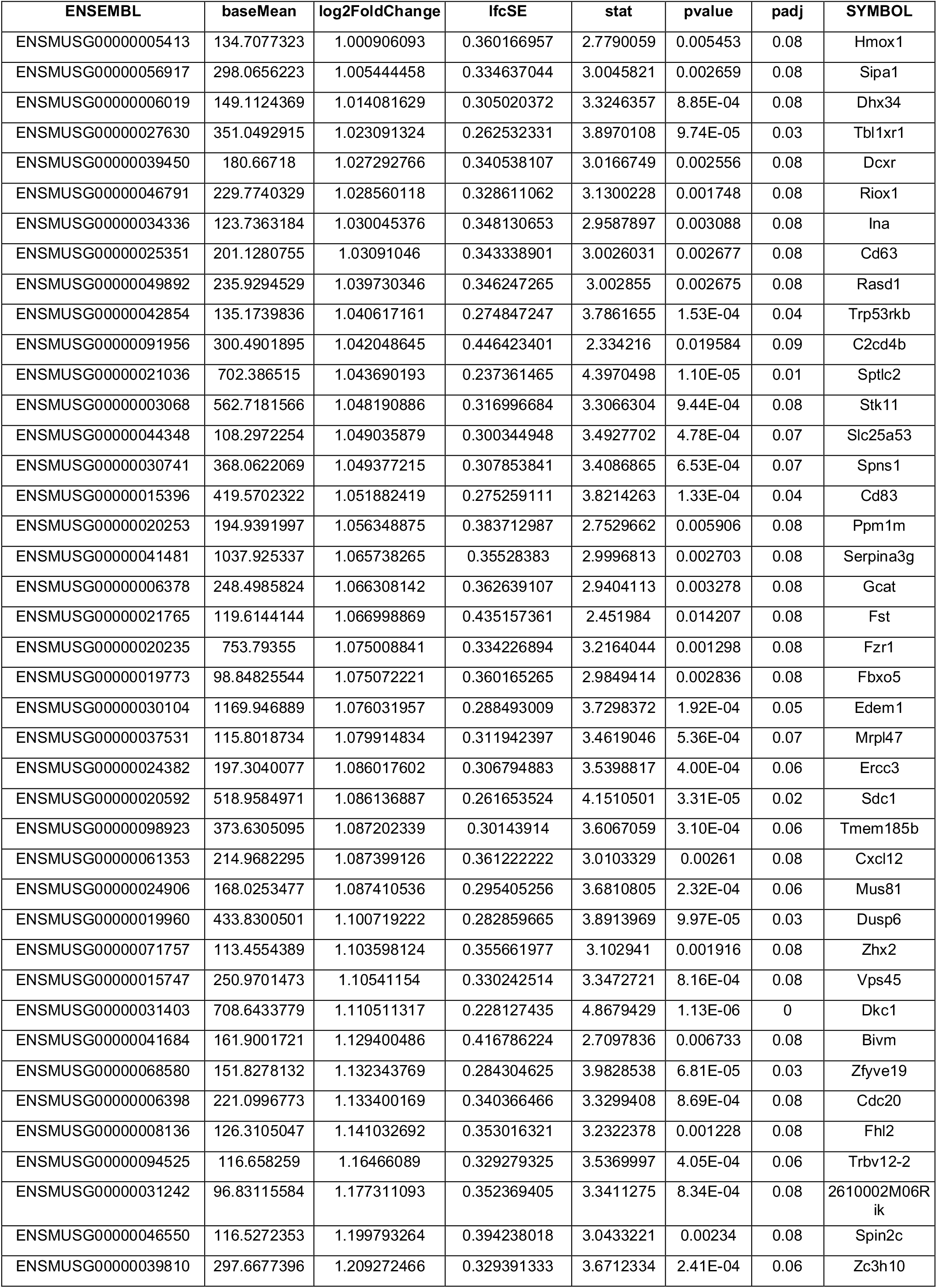
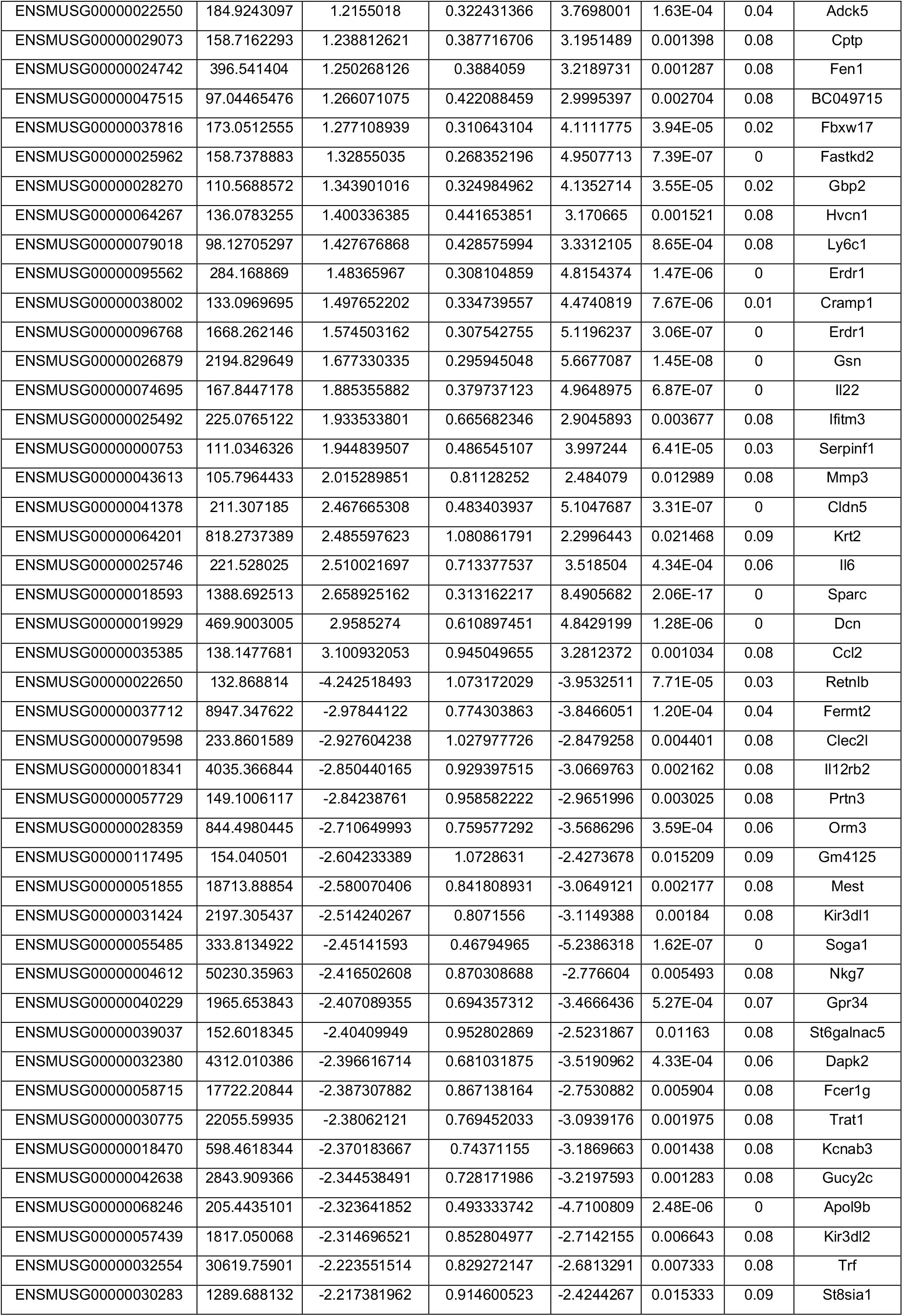
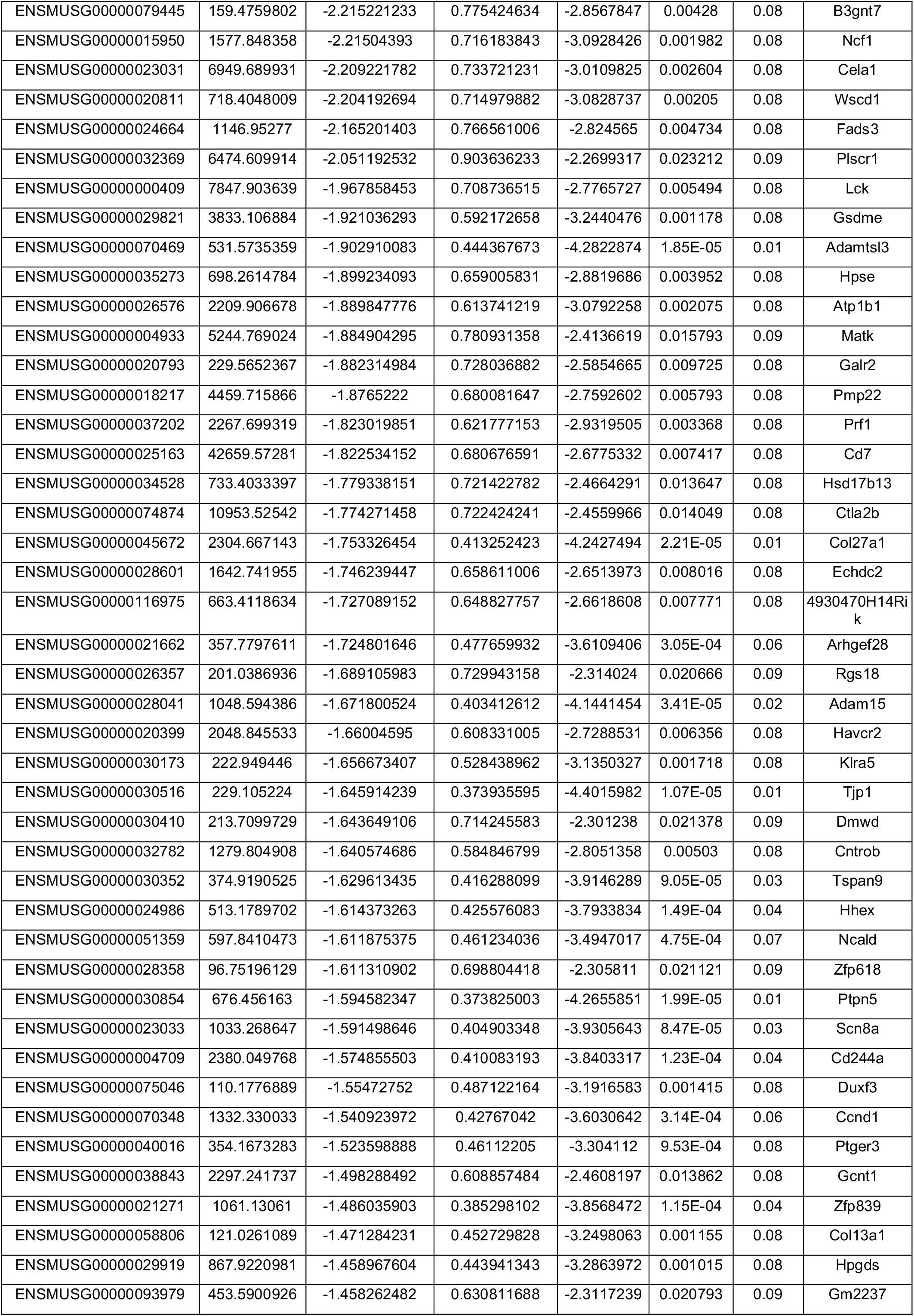
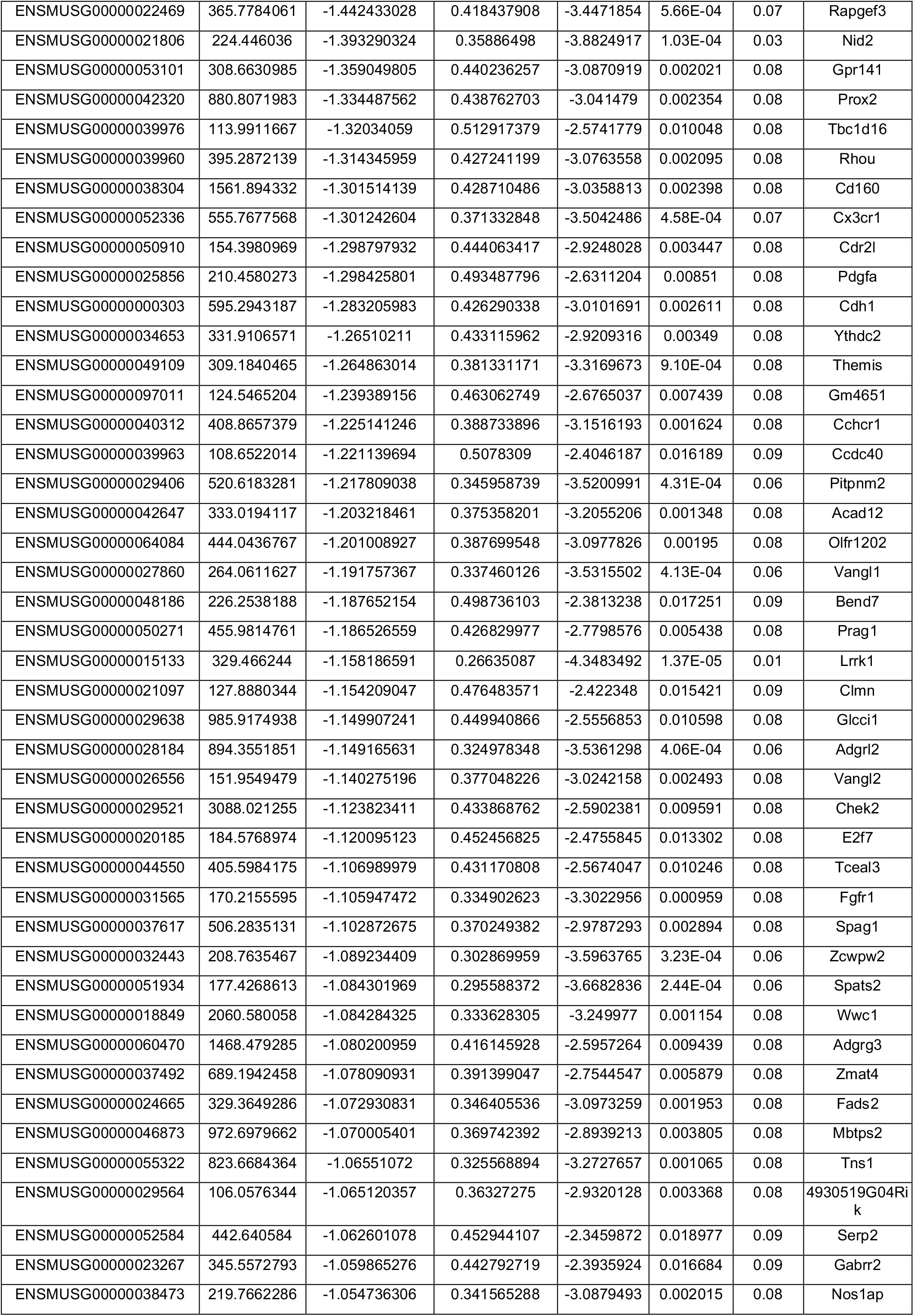
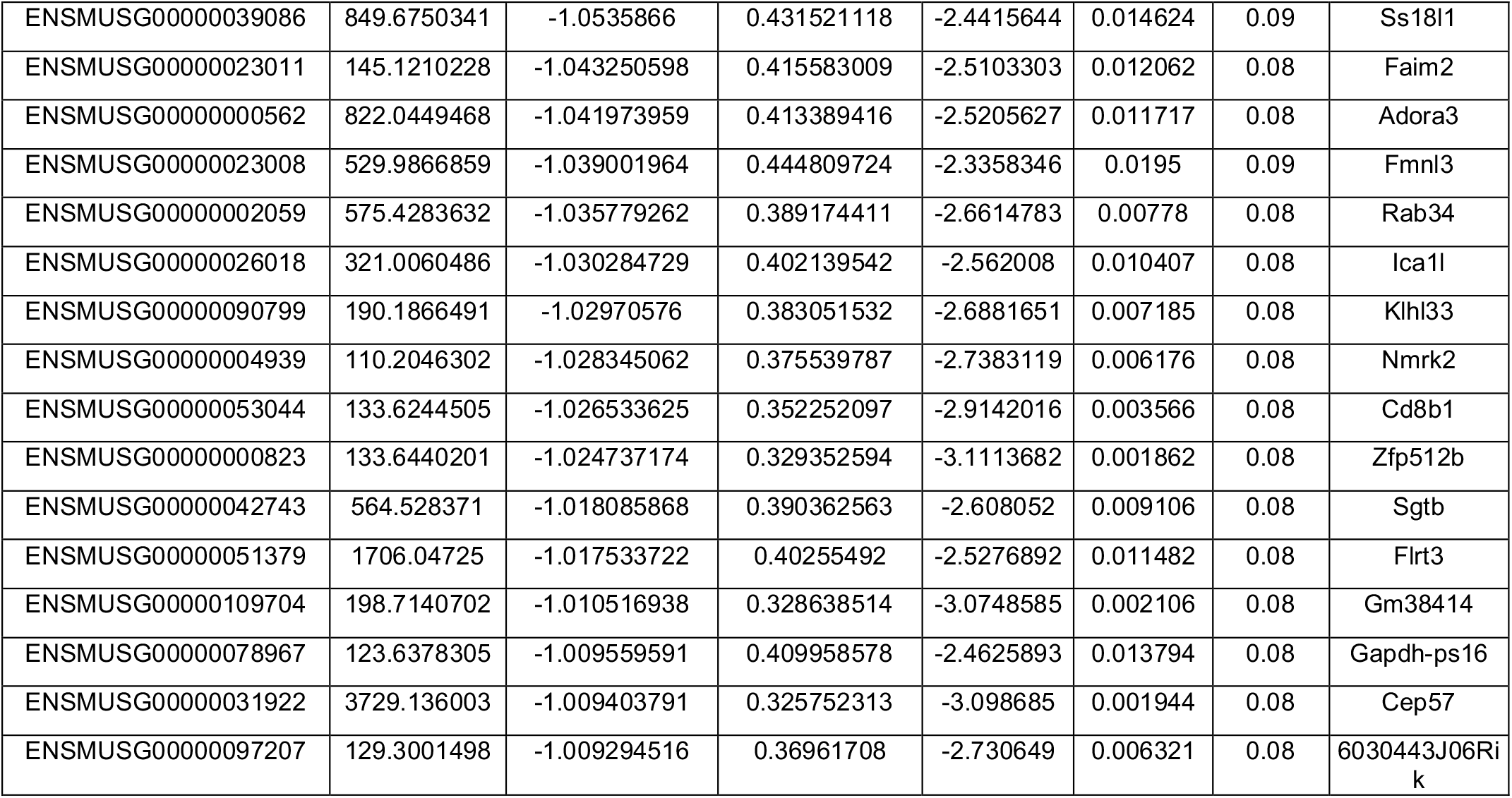

## Notes

### Competing Interest Statement

The authors have declared no competing interest.

### Summary of Updates

We added cell count data to most of the figures. We generated new data using intravenous CD45 labelling, showing that percutaneous, but not intravenous, N. brasiliensis infection induces an increase in interstitial γδ T cells in the lungs. We repeated key experiments to increase the number of replicates By using a temporal T Cell Receptor signalling reporter mouse line and cytokine blockade we defined the mechanism by which dermal γδ T cells are activated during N. brasiliensis infection.

## References

1. H. Veiga-Fernandes and A. A. Freitas. The s(c)ensory immune system theory. Trends Immunol, 38(10): 777–788, 2017. doi: 10.1016/j.it.2017.02.007.

2. P. Matzinger and T. Kamala. Tissue-based class control: the other side of tolerance. Nat Rev Immunol, 11(3): 221–30, 2011. doi: 10.1038/nri2940.

3. J. E. Allen, T. E. Sutherland, and D. Ruckerl. Il-17 and neutrophils: unexpected players in the type 2 immune response. Curr Opin Immunol, 34: 99–106, 2015. doi: 10.1016/j.coi.2015.03.001.

4. R. M. Maizels and W. C. Gause. Targeting helminths: The expanding world of type 2 immune effector mechanisms. J Exp Med, 220(10), 2023. doi: 10.1084/jem.20221381.

5. W. C. Gause, T. A. Wynn, and J. E. Allen. Type 2 immunity and wound healing: evolutionary refinement of adaptive immunity by helminths. Nat Rev Immunol, 13(8): 607–14, 2013. doi: 10.1038/nri3476.

6. J. E. Allen. Il-4 and il-13: Regulators and effectors of wound repair. Annu Rev Immunol, 41: 229–254, 2023. doi: 10.1146/annurev-immunol-101921-041206.

7. T. Bouchery, B. Volpe, K. Shah, L. Lebon, K. Filbey, G. LeGros, and N. Harris. The study of host immune responses elicited by the model murine hookworms nippostrongylus brasiliensis and heligmosomoides polygyrus. Curr Protoc Mouse Biol, 7(4): 236–286, 2017. doi: 10.1002/cpmo.34.

8. Jr. Urban, J. F., N. Noben-Trauth, D. D. Donaldson, K. B. Madden, S. C. Morris, M. Collins, and F. D. Finkelman. Il-13, il-4ralpha, and stat6 are required for the expulsion of the gastrointestinal nematode parasite nippostrongylus brasiliensis. Immunity, 8(2): 255–64, 1998. doi: 10.1016/s1074-7613(00)80477-x.

9. J. Ajendra, A. L. Chenery, J. E. Parkinson, B. H. K. Chan, S. Pearson, S. A. P. Colombo, L. Boon, R. K. Grencis, T. E. Sutherland, and J. E. Allen. Il-17a both initiates, via ifngamma suppression, and limits the pulmonary type-2 immune response to nematode infection. Mucosal Immunol, 13(6): 958–968, 2020. doi: 10.1038/s41385-020-0318-2.

10. T. E. Sutherland, N. Logan, D. Ruckerl, A. A. Humbles, S. M. Allan, V. Papayannopoulos, B. Stockinger, R. M. Maizels, and J. E. Allen. Chitinase-like proteins promote il-17-mediated neutrophilia in a tradeoff between nematode killing and host damage. Nat Immunol, 15(12): 1116–25, 2014. doi: 10.1038/ni.3023.

11. J. Ajendra, P. H. Papotto, J. E. Parkinson, R. J. Dodd, A. L. Bombeiro, S. Pearson, B. H. K. Chan, J. C. Ribot, H. J. McSorley, T. E. Sutherland, and J. E. Allen. The il-17a-neutrophil axis promotes epithelial cell il-33 production during nematode lung migration. Mucosal Immunol, 16(6): 767–775, 2023. doi: 10.1016/j.mucimm.2023.09.006.

12. F. Chen, W. Wu, L. Jin, A. Millman, M. Palma, D. W. El-Naccache, K. E. Lothstein, C. Dong, K. L. Edelblum, and W. C. Gause. B cells produce the tissue-protective protein relmalpha during helminth infection, which inhibits il-17 expression and limits emphysema. Cell Rep, 25(10):2775–2783 e3, 2018. doi: 10.1016/j.celrep.2018.11.038.

13. D. A. Ferrick, M. D. Schrenzel, T. Mulvania, B. Hsieh, W. G. Ferlin, and H. Lepper. Differential production of interferon-gamma and interleukin-4 in response to th1- and th2-stimulating pathogens by gamma delta t cells in vivo. Nature, 373(6511): 255–7, 1995. doi: 10.1038/373255a0.

14. J. C. Ribot, A. deBarros, D. J. Pang, J. F. Neves, V. Peperzak, S. J. Roberts, M. Girardi, J. Borst, A. C. Hayday, D. J. Pennington, and B. Silva-Santos. Cd27 is a thymic determinant of the balance between interferon-gamma- and interleukin 17-producing gammadelta t cell subsets. Nat Immunol, 10(4): 427–36, 2009. doi: 10.1038/ni.1717.

15. P. H. Papotto, J. C. Ribot, and B. Silva-Santos. Il-17(+) gammadelta t cells as kick-starters of inflammation. Nat Immunol, 18(6): 604–611, 2017. doi: 10.1038/ni.3726.

16. N. Sumaria, B. Roediger, L. G. Ng, J. Qin, R. Pinto, L. L. Cavanagh, E. Shklovskaya, B. Fazekas de St Groth, J. A. Triccas, and W. Weninger. Cutaneous immunosurveillance by self-renewing dermal gammadelta t cells. J Exp Med, 208(3): 505–18, 2011. doi: 10.1084/jem.20101824.

17. E. E. Gray, K. Suzuki, and J. G. Cyster. Cutting edge: Identification of a motile il-17-producing gammadelta t cell population in the dermis. J Immunol, 186(11): 6091–5, 2011. doi: 10.4049/jimmunol.1100427.

18. C. Benakis, D. Brea, S. Caballero, G. Faraco, J. Moore, M. Murphy, G. Sita, G. Racchumi, L. Ling, E. G. Pamer, C. Iadecola, and J. Anrather. Commensal microbiota affects ischemic stroke outcome by regulating intestinal gammadelta t cells. Nat Med, 22(5): 516–23, 2016. doi: 10.1038/nm.4068.

19. P. H. Papotto, B. Yilmaz, and B. Silva-Santos. Crosstalk between gammadelta t cells and the microbiota. Nat Microbiol, 6(9): 1110–1117, 2021. doi: 10.1038/s41564-021-00948-2.

20. D. L. Lee. Penetration of mammalian skin by the infective larva of nippostrongylus brasiliensis. Parasitology, 65(3): 499–505, 1972. doi: 10.1017/s0031182000044115.

21. L. M. Connor, S. C. Tang, E. Cognard, S. Ochiai, K. L. Hilligan, S. I. Old, C. Pellefigues, R. F. White, D. Patel, A. A. Smith, D. A. Eccles, O. Lamiable, M. J. McConnell, and F. Ronchese. Th2 responses are primed by skin dendritic cells with distinct transcriptional profiles. J Exp Med, 214(1): 125–142, 2017. doi: 10.1084/jem.20160470.

22. C. Pellefigues, S. C. Tang, A. Schmidt, R. F. White, O. Lamiable, L. M. Connor, C. Ruedl, J. Dobrucki, G. Le Gros, and F. Ronchese. Toll-like receptor 4, but not neutrophil extracellular traps, promote ifn type i expression to enhance th2 responses to nippostrongylus brasiliensis. Front Immunol, 8:1575, 2017. doi: 10.3389/fimmu.2017.01575.

23. H. Nagashima, T. Mahlakoiv, H. Y. Shih, F. P. Davis, F. Meylan, Y. Huang, O. J. Harrison, C. Yao, Y. Mikami, Jr. Urban, J. F., K. M. Caron, Y. Belkaid, Y. Kanno, D. Artis, and J. J. O’Shea. Neuropeptide cgrp limits group 2 innate lymphoid cell responses and constrains type 2 inflammation. Immunity, 51(4):682–695 e6, 2019. doi: 10.1016/j.immuni.2019.06.009.

24. F. Chen, Z. Liu, W. Wu, C. Rozo, S. Bowdridge, A. Millman, N. Van Rooijen, Jr. Urban, J. F., T. A. Wynn, and W. C. Gause. An essential role for th2-type responses in limiting acute tissue damage during experimental helminth infection. Nat Med, 18(2): 260–6, 2012. doi: 10.1038/nm.2628.

25. K. G. Anderson, K. Mayer-Barber, H. Sung, L. Beura, B. R. James, J. J. Taylor, L. Qunaj, T. S. Griffith, V. Vezys, D. L. Barber, and D. Masopust. Intravascular staining for discrimination of vascular and tissue leukocytes. Nat Protoc, 9(1): 209–22, 2014. ISSN 1750-2799 (Electronic) 1754-2189 (Print) 1750-2799 (Linking). doi: 10.1038/nprot.2014.005.

26. B. Liu, L. Bao, L. Wang, F. Li, M. Wen, H. Li, W. Deng, X. Zhang, and B. Cao. Anti-ifn-gamma therapy alleviates acute lung injury induced by severe influenza a (h1n1) pdm09 infection in mice. J Microbiol Immunol Infect, 54(3): 396–403, 2021. doi: 10.1016/j.jmii.2019.07.009.

27. R. Wiesheu, S. C. Edwards, A. Hedley, H. Hall, M. Tosolini, M. G. F. Fares da Silva, N. Sumaria, S. M. Castenmiller, L. Wardak, Y. Optaczy, A. Lynn, D. G. Hill, A. J. Hayes, J. Hay, A. Kilbey, R. Shaw, D. Whyte, P. J. Walsh, A. M. Michie, G. J. Graham, A. Manoharan, C. Halsey, K. Blyth, M. C. Wolkers, C. Miller, D. J. Pennington, G. W. Jones, J. J. Fournie, V. Bekiaris, and S. B. Coffelt. Il-27 maintains cytotoxic ly6c(+) gammadelta t cells that arise from immature precursors. EMBO J, 43(14): 2878–2907, 2024. doi: 10.1038/s44318-024-00133-1.

28. L. Mazzurana, P. Czarnewski, V. Jonsson, L. Wigge, M. Ringner, T. C. Williams, A. Ravindran, A. K. Bjorklund, J. Safholm, G. Nilsson, S. E. Dahlen, A. C. Orre, M. Al-Ameri, C. Hoog, C. Hedin, S. Szczegielniak, S. Almer, and J. Mjosberg. Tissue-specific transcriptional imprinting and heterogeneity in human innate lymphoid cells revealed by full-length single-cell rna-sequencing. Cell Res, 31(5): 554–568, 2021. doi: 10.1038/s41422-020-00445-x.

29. A. P. McFarland, A. Yalin, S. Y. Wang, V. S. Cortez, T. Landsberger, R. Sudan, V. Peng, H. L. Miller, B. Ricci, E. David, R. Faccio, I. Amit, and M. Colonna. Multi-tissue single-cell analysis deconstructs the complex programs of mouse natural killer and type 1 innate lymphoid cells in tissues and circulation. Immunity, 54(6):1320–1337 e4, 2021. doi: 10.1016/j.immuni.2021.03.024.

30. T. Bouchery, M. Moyat, J. Sotillo, S. Silverstein, B. Volpe, G. Coakley, T. D. Tsourouktsoglou, L. Becker, K. Shah, M. Kulagin, R. Guiet, M. Camberis, A. Schmidt, A. Seitz, P. Giacomin, G. Le Gros, V. Papayannopoulos, A. Loukas, and N. L. Harris. Hookworms evade host immunity by secreting a deoxyribonuclease to degrade neutrophil extracellular traps. Cell Host Microbe, 27(2):277–289 e6, 2020. doi: 10.1016/j.chom.2020.01.011.

31. F. Ramirez-Valle, E. E. Gray, and J. G. Cyster. Inflammation induces dermal vgamma4+ gammadeltat17 memory-like cells that travel to distant skin and accelerate secondary il-17-driven responses. Proc Natl Acad Sci U S A, 112(26): 8046–51, 2015. doi: 10.1073/pnas.1508990112.

32. D. R. McKenzie, E. E. Kara, C. R. Bastow, T. S. Tyllis, K. A. Fenix, C. E. Gregor, J. J. Wilson, R. Babb, J. C. Paton, A. Kallies, S. L. Nutt, A. Brustle, M. Mack, I. Comerford, and S. R. McColl. Il-17-producing gammadelta t cells switch migratory patterns between resting and activated states. Nat Commun, 8:15632, 2017. doi: 10.1038/ncomms15632.

33. G. M. Halliday, D. L. Damian, S. Rana, and S. N. Byrne. The suppressive effects of ultraviolet radiation on immunity in the skin and internal organs: implications for autoimmunity. J Dermatol Sci, 66(3): 176–82, 2012. ISSN 1873-569X (Electronic) 0923-1811 (Linking). doi: 10.1016/j.jdermsci.2011.12.009.

34. M. Petkova, M. Kraft, S. Stritt, I. Martinez-Corral, H. Ortsater, M. Vanlandewijck, B. Jakic, E. Baselga, S. D. Castillo, M. Graupera, C. Betsholtz, and T. Makinen. Immune-interacting lymphatic endothelial subtype at capillary terminals drives lymphatic malformation. J Exp Med, 220(4), 2023. ISSN 1540-9538 (Electronic) 0022-1007 (Print) 0022-1007 (Linking). doi: 10.1084/jem.20220741.

35. T. A. E. Elliot, E. K. Jennings, D. A. J. Lecky, S. Rouvray, G. M. Mackie, L. Scarfe, L. Sheriff, M. Ono, K. M. Maslowski, and D. Bending. Nur77-tempo mice reveal t cell steady state antigen recognition. Discov Immunol, 1(1):kyac009, 2022. ISSN 2754-2483 (Electronic) 2754-2483 (Linking). doi: 10.1093/discim/kyac009.

36. Y. Cai, X. Shen, C. Ding, C. Qi, K. Li, X. Li, V. R. Jala, H. G. Zhang, T. Wang, J. Zheng, and J. Yan. Pivotal role of dermal il-17-producing gammadelta t cells in skin inflammation. Immunity, 35(4): 596–610, 2011. ISSN 1097-4180 (Electronic) 1074-7613 (Print) 1074-7613 (Linking). doi: 10.1016/j.immuni.2011.08.001.

37. P. H. Papotto, N. Goncalves-Sousa, N. Schmolka, A. Iseppon, S. Mensurado, B. Stockinger, J. C. Ribot, and B. Silva-Santos. Il-23 drives differentiation of peripheral gammadelta17 t cells from adult bone marrow-derived precursors. EMBO Rep, 18(11): 1957–1967, 2017. ISSN 1469-3178 (Electronic) 1469-221X (Print) 1469-221X (Linking). doi: 10.15252/embr.201744200.

38. S. J. Van Dyken, A. Mohapatra, J. C. Nussbaum, A. B. Molofsky, E. E. Thornton, S. F. Ziegler, A. N. McKenzie, M. F. Krummel, H. E. Liang, and R. M. Locksley. Chitin activates parallel immune modules that direct distinct inflammatory responses via innate lymphoid type 2 and gammadelta t cells. Immunity, 40(3): 414–24, 2014. ISSN 1097-4180 (Electronic) 1074-7613 (Print) 1074-7613 (Linking). doi: 10.1016/j.immuni.2014.02.003.

39. J. M. Inclan-Rico, C. M. Napuri, C. Lin, L. Y. Hung, A. A. Ferguson, X. Liu, Q. Wu, C. F. Pastore, A. Stephenson, U. M. Femoe, F. Musaigwa, H. L. Rossi, B. D. Freedman, D. R. Reed, T. Machacek, P. Horak, I. Abdus-Saboor, W. Luo, and D. R. Herbert. Mrgpra3 neurons drive cutaneous immunity against helminths through selective control of myeloid-derived il-33. Nat Immunol, 25(11): 2068–2084, 2024. doi: 10.1038/s41590-024-01982-y.

40. J. M. Inclan-Rico, A. Stephenson, C. M. Napuri, H. L. Rossi, L. Y. Hung, C. F. Pastore, W. Luo, and D. R. Herbert. Trpv1+ neurons promote cutaneous immunity against schistosoma mansoni. J Immunol, 2025. ISSN 1550-6606 (Electronic) 0022-1767 (Linking). doi: 10.1093/jimmun/vkaf141.

41. T. Hartwig, S. Pantelyushin, A. L. Croxford, P. Kulig, and B. Becher. Dermal il-17-producing gammadelta t cells establish long-lived memory in the skin. Eur J Immunol, 45(11): 3022–33, 2015. doi: 10.1002/eji.201545883.

42. K. de Ruiter, S. P. Jochems, D. L. Tahapary, K. A. Stam, M. Konig, V. van Unen, S. Laban, T. Hollt, M. Mbow, B. P. F. Lelieveldt, F. Koning, E. Sartono, J. W. A. Smit, T. Supali, and M. Yazdanbakhsh. Helminth infections drive heterogeneity in human type 2 and regulatory cells. Sci Transl Med, 12(524), 2020. ISSN 1946-6242 (Electronic) 1946-6234 (Linking). doi: 10.1126/scitranslmed.aaw3703.

43. B. Xie, M. Wang, X. Zhang, Y. Zhang, H. Qi, H. Liu, Y. Wu, X. Wen, X. Chen, M. Han, D. Xu, X. Sun, X. Zhang, X. Zhao, Y. Shang, S. Yuan, and J. Zhang. Gut-derived memory gammadelta t17 cells exacerbate sepsis-induced acute lung injury in mice. Nat Commun, 15(1): 6737, 2024. doi: 10.1038/s41467-024-51209-9.

44. C. T. Ng, L. Y. Fong, M. R. Sulaiman, M. A. Moklas, Y. K. Yong, M. N. Hakim, and Z. Ahmad. Interferon-gamma increases endothelial permeability by causing activation of p38 map kinase and actin cytoskeleton alteration. J Interferon Cytokine Res, 35(7): 513–22, 2015. doi: 10.1089/jir.2014.0188.

45. V. Langer, E. Vivi, D. Regensburger, T. H. Winkler, M. J. Waldner, T. Rath, B. Schmid, L. Skottke, S. Lee, N. L. Jeon, T. Wohlfahrt, V. Kramer, P. Tripal, M. Schumann, S. Kersting, C. Handtrack, C. I. Geppert, K. Suchowski, R. H. Adams, C. Becker, A. Ramming, E. Naschberger, N. Britzen-Laurent, and M. Sturzl. Ifn-gamma drives inflammatory bowel disease pathogenesis through ve-cadherin-directed vascular barrier disruption. J Clin Invest, 129(11): 4691–4707, 2019. doi: 10.1172/JCI124884.

46. D. A. Hill and J. M. Spergel. The atopic march: Critical evidence and clinical relevance. Ann Allergy Asthma Immunol, 120(2): 131–137, 2018. doi: 10.1016/j.anai.2017.10.037.

47. H. M. Lee, A. Fleige, R. Forman, S. Cho, A. A. Khan, L. L. Lin, D. T. Nguyen, A. O’Hara-Hall, Z. Yin, C. A. Hunter, W. Muller, and L. F. Lu. Ifngamma signaling endows dcs with the capacity to control type i inflammation during parasitic infection through promoting t-bet+ regulatory t cells. PLoS Pathog, 11(2):e1004635, 2015. doi: 10.1371/journal.ppat.1004635.

48. M. Camberis, G. Le Gros, and Jr. Urban, J. Animal model of nippostrongylus brasiliensis and heligmosomoides polygyrus. Curr Protoc Immunol, Chapter 19:Unit 19 12, 2003. doi: 10.1002/0471142735.im1912s55.

49. E. Dahlberg. Estimation of the blood contamination of tissue extracts. Anal Biochem, 130 (1):108–13, 1983. doi: 10.1016/0003-2697(83)90656-5.

50. R. A. Amezquita, A. T. L. Lun, E. Becht, V. J. Carey, L. N. Carpp, L. Geistlinger, F. Marini, K. Rue-Albrecht, D. Risso, C. Soneson, L. Waldron, H. Pages, M. L. Smith, W. Huber, M. Morgan, R. Gottardo, and S. C. Hicks. Orchestrating single-cell analysis with bioconductor. Nat Methods, 17(2): 137–145, 2020. doi: 10.1038/s41592-019-0654-x.

51. I. Dean, B. C. Kennedy, Z. Li, F. Berditchevski, and D. R. Withers. Protocol for transcutaneous tumor photolabeling to track immune cells in vivo using kaede mice. STAR Protoc, 5(2): 102956, 2024. ISSN 2666-1667 (Electronic) 2666-1667 (Linking). doi: 10.1016/j.xpro.2024.102956.

